# Co‑cultivation of the anaerobic fungus *Caecomyces churrovis* with *Methanobacterium bryantii* enhances transcription of carbohydrate binding modules

**DOI:** 10.1101/2021.07.09.451685

**Authors:** Jennifer L. Brown, Candice L. Swift, Stephen Mondo, Susanna Seppala, Asaf Salamov, Vasanth Singan, Bernard Henrissat, John K. Henske, Samantha Lee, Guifen He, Mi Yan, Kerrie Barry, Igor V. Grigoriev, Michelle A. O’Malley

## Abstract

Anaerobic fungi and methanogenic archaea are two classes of microorganisms found in the rumen microbiome that metabolically interact during lignocellulose breakdown. Here, stable synthetic co-cultures of the anaerobic fungus *Caecomyces churrovis* and the methanogen *Methanobacterium bryantii* (not native to the rumen) were formed, demonstrating that microbes from different environments can be paired based on metabolic ties. Transcriptional and metabolic changes induced by methanogen co-culture were evaluated in *C. churrovis* across a variety of substrates to identify mechanisms that impact biomass breakdown and sugar uptake. A high-quality genome of *C. churrovis* was obtained and annotated, which is the first sequenced genome of a non-rhizoid forming anaerobic fungus. *C. churrovis* possess an abundance of CAZymes and carbohydrate binding modules and, in agreement with previous studies of early-diverging fungal lineages, N6-methyldeoxyadenine (6mA) was associated with transcriptionally active genes. Co-culture with the methanogen increased overall transcription of CAZymes, carbohydrate binding modules, and dockerin domains in co-cultures grown on both lignocellulose and cellulose and caused upregulation of genes coding associated enzymatic machinery including carbohydrate binding modules in family 18 and dockerin domains across multiple growth substrates relative to *C. churrovis* monoculture. Two other fungal strains grown on a reed canary grass substrate in co-culture with the same methanogen also exhibited high log2fold change values for upregulation of genes encoding carbohydrate binding modules in families 1 and 18. Transcriptional upregulation indicated that co-culture of the *C. churrovis* strain with a methanogen may enhance pyruvate formate lyase (PFL) function for growth on xylan and fructose and production of bottleneck enzymes in sugar utilization pathways, further supporting the hypothesis that co-culture with a methanogen may enhance certain fungal metabolic functions. Upregulation of CBM18 may play a role in fungal-methanogen physical associations and fungal cell wall development and remodeling.

## Introduction

Anaerobic fungi are efficient degraders of recalcitrant lignocellulosic biomass that are found in the guts of herbivores. The high number of CAZymes (carbohydrate active enzymes) that anaerobic fungi produce has driven efforts to collect genomic and transcriptomic data for a variety of emerging anaerobic fungal species ^1–5^. Gut fungi function within a community of biomass-degrading bacteria, protozoa, and methanogenic archaea linked by complex metabolic interactions and functional redundancy.^6^ Isolating individual members of these natural consortia is one approach to develop a more detailed understanding of microbial interactions, which can then be used to design optimized consortia for biotechnological applications to break down lignocellulose-rich waste. These microbes can be selected through “top-down” enrichment techniques such as serial cultivation or antibiotic treatment to isolate syntrophic pairs of fungi and methanogens from naturally-occurring consortia. Alternatively, communities can be formed using “bottom up” methods mixing separate axenic cultures of these microbes to create synthetic pairings linked by metabolic dependency.^6–8^

Fungal-methanogen co-cultures have been extensively studied due to the mutually beneficial relationship between the two organisms resulting from their complementary metabolism. – fungi produce hydrogen (H_2_) as an unwelcome byproduct of their own metabolism, which methanogens use in the biosynthesis and release of methane ^8–14^ Many previous studies report that co-cultivation of anaerobic fungi with methanogens can enhance biomass breakdown, but the metabolic mechanisms responsible for this outcome are unclear and not uniformly reproducible.^15–19^ For example, a recent study concluded that the removal of fungal metabolites by methanogens does not increase the rate of gas production or the rate of substrate deconstruction by a synthetic community of fungi and methanogens relative to fungal monocultures.^8^ It has also been hypothesized that co-cultivation of fungi and methanogens results in increased sugar utilization and flux through the fungal hydrogenosome through increased transport and carbon conversion.^14,20^ Additionally, we recently reported that *M. bryantii* enhances the transcription of genes encoding ABC transporters, MFS transporters and G-protein coupled receptors (GPCRs) in the fungus *Anaeromyces robustus*, indicating that co-cultivation may increase the rate of sugar utilization through the increased expression of sugar transporters.^9^ Although many studies have been conducted to determine how co-cultivation with methanogens affects fungal metabolism and biomass breakdown, none have characterized transcriptional and metabolic outcomes across a variety of relevant substrates, which is critical to detangling competing effects of substrate response.^9,10^

Here, we present the first genome of an anaerobic non-rhizoid forming fungus of the *Caecomyces* genus, and further examine its transcriptional response to the presence of methanogens in multiple synthetic co-cultures supported on lignocellulose, hemicellulose, cellulose, and sugars. *Caecomyces churrovis* lacks the extensive rhizoid network formed by other previously sequenced anaerobic gut fungi to aid in biomass breakdown. Improvements in long-read sequencing technologies enabled assembly and annotation of CAZymes and associated cellular machinery despite the complex fungal physiology, unknown ploidy, AT-content, and repeat-richness. By combining RNA-seq with growth and chemical data, we determine how the fungus responds to co-cultivation with a non-native methanogen in synthetic co-culture. While other studies have examined global transcriptomic response and CAZyme regulation in anaerobic fungi cultivated with methanogens on a single substrate, none to date have explored regulation across a range of substrates or differences occurring in transcriptional regulation between multiple fungal strains on the same substrate.^9,10^ Through a combination of genomic, transcriptomic, and metabolomic data we found that the *Caecomyces churrovis* genome possesses an abundance of both CAZymes and carbohydrate binding modules as shown in Figure 1. Co-culture of *C. churrovis* with a non-native methanogen enhances transcription of gene sets associated with fungal substrate binding and fungal-methanogen interactions such as carbohydrate binding modules in families 1 and 18, pyruvate formate lyase (PFL) function in the cytosol or possibly the hydrogenosome, and enzymes that are potential bottlenecks for sugar utilization in fungi across multiple substrates. Overall, understanding how methanogen co-culture influences the fibrolytic and metabolic behavior of anaerobic fungi aids in the design of new strategies for conversion of lignocellulose to fermentable sugars and value-added products, as well as the genetic mechanisms that underpin fungal-methanogen interactions.

**Figure 1.**
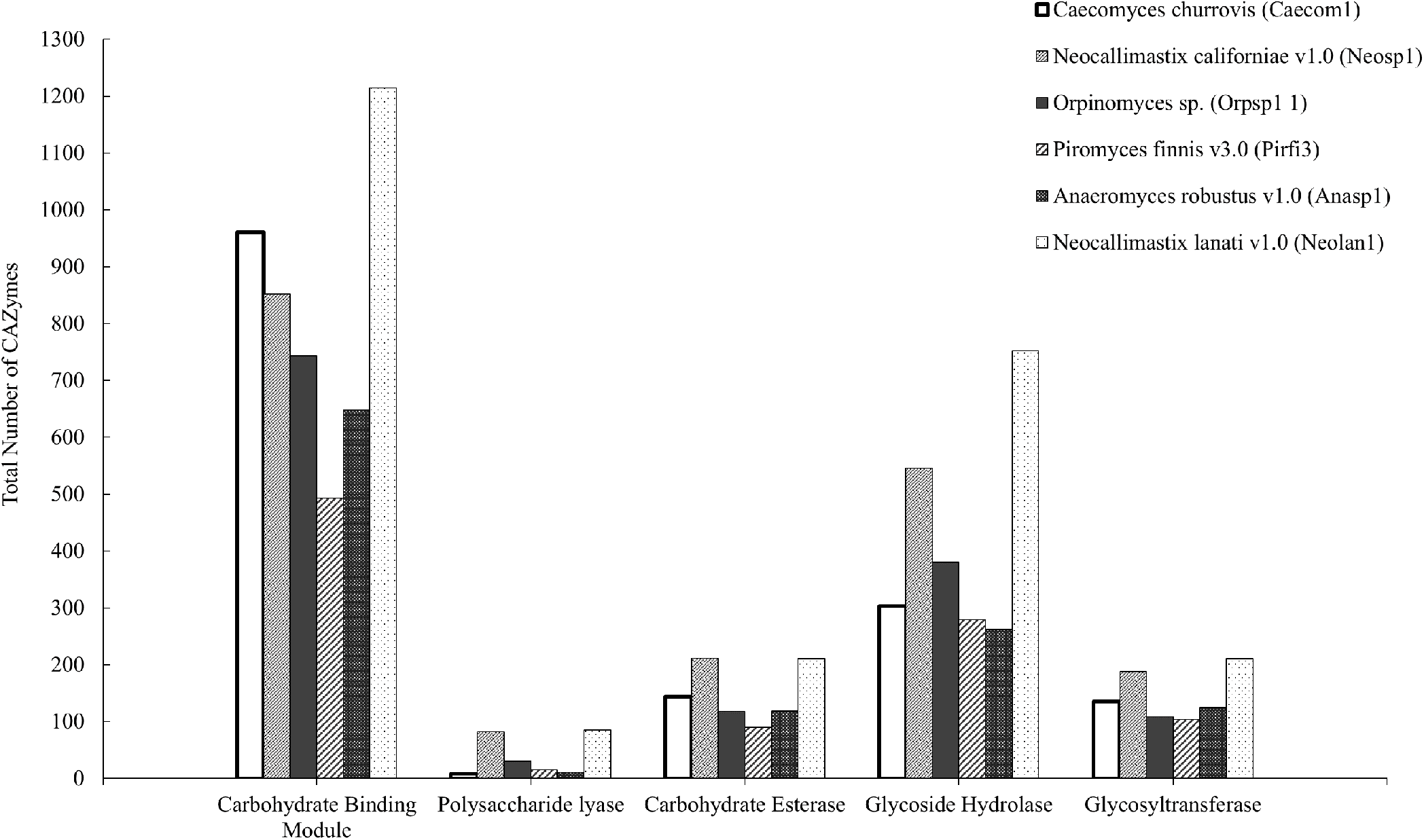
Number of different types of CAZyme domains in six sequenced anaerobic fungi. *C. churrovis* has the highest number of domains annotated as carbohydrate-binding modules compared to most other sequenced anaerobic fungi. Annotation data for these strains can be found at https://mycocosm.jgi.doe.gov.

## Methods

### Growing and harvesting cultures for RNA extractions

Anaerobic serum bottles containing 80 mL of modified medium C (“MC-”) with 0.8 mL 100 × vitamin solution and 0.8 g reed canary grass were inoculated with cultures of *C. churrovis* and *M. bryantii*: 1.0 mL of *C. churrovis* or a combination of 1.0 mL of *C. churrovis* and 1.0 mL of *M. bryantii* (routine cultures were cultivated as described previously by Swift, et al.).^9^ The fungal and methanogen co-cultures and fungal monocultures were grown anaerobically at 39°C in Hungate tubes filled with 9.0 mL of autoclaved modified medium C^21^ (“MC-”), containing 1.25 g/L yeast extract, 5 g/L Bacto™ Casitone, and 7.5 vol% clarified rumen fluid, with either 0.1 g of milled reed canary grass, 0.1 g Avicel, 0.1 g Xylan, 0.5 ml of a 0.1 g/ml sterile filtered glucose stock solution, or 0.1 g/ml of a sterile filtered fructose stock solution as the growth substrate, and supplemented with vitamin solution post-autoclaving.^22^ Pressure production was used as a proxy for fungal growth, as described previously.^23^ Daily pressure measurements were taken using a probe pressure transducer to determine when the cultures reached the mid-log growth phase, based upon previous pressure growth curves measured to stationary phase growth. Upon reaching mid-log growth phase, cultures were harvested and stored for later RNA extraction. After sampling the headspace gas of the culture to determine end-point methane and hydrogen concentrations for monocultures and co-cultures, a volume of 1.2 ml of the culture supernatant was pipetted off of the top of the culture and stored at −20°C for later HPLC analysis. The remainder of the culture was transferred to a 15 ml falcon tube and spun down at 10,000 g and 4°C for 6 minutes. The remaining supernatant was then decanted or pipetted off depending upon the integrity of the remaining cell pellet and replaced with 1 ml of RNA-later and mixed by pipetting. Samples were then stored at −80C until extraction.

### Measuring Hydrogen and Methane Production

End-point methane and hydrogen measurements for both monocultures and co-cultures were taken from the headspace of the culture tubes before harvesting the cultures. Daily measurements and sampling were performed to monitor the growth of the co-cultures and monocultures in the following order. First the pressure in each sample was measured using a pressure transducer,^24^ and the headspace composition was measured on a gas chromatograph (GC)-pulsed, discharge helium ionization detector (Thermo Fisher Scientific TRACE 1300).^25^ Finally, the headspace pressure of the sample was vented return the headspace to atmospheric pressure. The total moles of headspace gas were calculated using the ideal gas law. Gas concentrations for H_2_ and methane were calculated using an external standard calibration method. The gas concentration could then be multiplied by the number of moles present both before and after the pressure sampling in order to determine the moles of H_2_ or methane produced. It was assumed that the amount of gas dissolved in the liquid media was negligible for these calculations.

### HPLC Analysis

Levels of volatile fatty acids present in the supernatant of both co-cultures and monocultures were measured using an Agilent1260 Infinity HPLC (Agilent). Samples were prepared by acidifying to 5 mM using sulfuric acid and subsequently incubating at room temperature for 5 minutes. Samples were then centrifuged for 5 minutes at 21,000g. The supernatant was syringe filtered into an HPLC vial (Eppendorf FA-45-24-11) using a 0.22 μm PVDF filter. Samples were analyzed on an Agilent 1260 Infinity high-performance liquid chromatography system (HPLC, Agilent, Santa Clara, CA) equipped with an auto-sampler unit (1260 ALS). Separation of formate, acetate, and lactate was achieved with a Bio-Rad Aminex® 87H Ion Exclusion Column for organic acids (Part No. 1250140, Bio-Rad, Hercules, CA) with a mobile phase of 5 mM sulfuric acid. In-house standards were prepared with MC- blank culture medium as a base and sodium formate (ACS Grade, Fisher Chemical S648500), sodium acetate (ACS Grade, Fisher Chemical S210500), L-lactic acid sodium (99%, extra pure, Acros Organics 439220100) at VFA concentrations of 0.1 and 1 g/L.

### Genome Sequencing and Annotation of Anaerobic Fungus *Caecomyces churrovis*

The *Caecomyces churrovis* genome was sequenced using the PacBio sequencing platform. >10kb fragments were size selected using Blue Pippin Size Selection, then 10 ug of genomic DNA was sheared to >10kb fragments using Covaris g-Tubes. The sheared DNA was treated with exonuclease to remove single-stranded ends and DNA damage repair mix followed by end repair and ligation of blunt adapters using SMRTbell Template Prep Kit 1.0 (Pacific Biosciences). The library was purified with AMPure PB beads and size selected with BluePippin (Sage Science) at >10 kb cutoff size. PacBio Sequencing primer was then annealed to the SMRTbell template library and sequencing polymerase was bound to them using Sequel Binding kit 2.0. The prepared SMRTbell template libraries were then sequenced on a Pacific Biosystems’ Sequel sequencer using v3 sequencing primer, 1M v2 SMRT cells, and Version 2.0 sequencing chemistry with 6 hour & 10 hour movie run times. 6mA modifications were detected using the PacBio SMRT analysis platform (pb_basemods package; smrtanalysis version: smrtlink/8.0.0.80529). 6mA modifications were then filtered and methylated genes were identified following the methods described in Mondo et al., 2017.^26^ The assembly was completed with Falcon (https://github.com/PacificBiosciences/FALCON) which generates better assemblies than competing methods likely due to an improvement in isolation of high molecular weight DNA and sequencing larger DNA fragments.^1^ While annotating fungal genomes present a challenge due to the lack of anaerobic fungal gene content in existing databases, the genome was annotated using the JGI Annotation Pipeline, which employs a variety of gene modelers to discover genes. In addition to homology-based modelers, ab-initio gene discovery tools and RNAseq based methods were used for annotation. Models were determined to be allelic if they were located in regions on smaller scaffolds that were > 95% identical at the nucleic acid level and > 50% of the smaller scaffold was covered by these regions.

### Extracting RNA from Experimental Samples

Samples were removed from storage at −80C and thawed on ice. After thawing, samples were spun down for 6 minutes at 4°C and 10,000 g and RNA later was removed. Cells were lysed for the reed canary grass and Avicel cultures using bead beating for 1 minute in 30 second intervals. Total RNA was extracted using the RNeasy Mini kit (QIAGEN) following the protocol for “Purification of Total RNA from Plant Cells and Tissues and Filamentous Fungi” including an on-column DNAse digest. An Agilent TapeStation was used to determine the quality of the sequenced RNA and Qubit High Sensitivity RNA Assay was used to determine concentrations.

### RNA Sequencing and Data Analysis

Stranded RNASeq library(s) were created and quantified by qPCR for both monoculture and co-culture samples. Stranded cDNA libraries were generated using the Illumina Truseq Stranded mRNA Library Prep kit. mRNA was purified from 1 ug of total RNA using magnetic beads containing poly-T oligos. mRNA was fragmented and reversed transcribed using random hexamers and SSII (Invitrogen) followed by second strand synthesis. The fragmented cDNA was treated with end-pair, A-tailing, adapter ligation, and 8 cycles of PCR. The prepared library was quantified using KAPA Biosystems’ next-generation sequencing library qPCR kit and run on a Roche LightCycler 480 real-time PCR instrument. The quantified library was then prepared for sequencing on the Illumina HiSeq sequencing platform utilizing a TruSeq paired-end cluster kit, v4. Sequencing of the flow cell was performed on the Illumina HiSeq 2500 sequencer using HiSeq TruSeq SBS sequencing kits, v4, following a 2×150 indexed run recipe. Sequencing was performed using an Illumina® Novaseq. The filtered reads from each library were aligned to the *Caecomyces churrovis* genome using HISAT2 version 2.1.0.^27^ Strand-specific coverage was generated using deepTools v3.1.^28^ Raw gene counts were generated using featureCounts, with only primary hits assigned to the reverse strand were included in the raw gene counts.^29^ Raw gene counts were used to evaluate the level of correlation between biological replicates using Pearson’s correlation and determine which replicates would be used in the DGE analysis. DESeq2 (version 1.18.1)^30^ was subsequently used to determine which genes were differentially expressed between pairs of conditions. The parameters used to call a gene DE between conditions were p-value < 0.05 and a log2fold change greater than 2. This log2fold change cutoff is more stringent than the typical cutoff used in previous studies to account for variation in undefined rumen fluid components across different batches of media. Raw gene counts, not normalized counts, were used for DGE analysis since DESeq2 uses its own internal normalization. Subsequent analysis was done using the filtered model gene catalog for *C. churrovis* provided for download on the Mycocosm website.

## Results and Discussion

### The Caecomyces churrovis genome encodes an abundance of CAZymes and carbohydrate binding modules

Anaerobic fungi are emerging platforms for hydrolysis of crude lignocellulose, as they produce powerful CAZymes and mechanically associate with and often penetrate plant cell walls.^5,31,32^ The first high quality genome of a non-rhizoid forming anaerobic fungus from the *Caecomyces* genera was sequenced with PacBio SMRT sequencing using high molecular weight DNA fragments, a method that is critical to high-quality genome assemblies for anaerobic fungi.^33–35^ Previously, we assembled a *de novo* transcriptome of *C. churrovis* by pooling RNA from batch cultures grown on glucose, cellobiose, cellulose, and reed canary grass, obtaining an inclusive set of expressed genes for these substrates.^5^ The acquisition of the *C. churrovi*s genome now enables more detailed investigation of genetic regulatory mechanisms, splicing, ploidy, and comparative genomics that cannot be accomplished with a sole transcriptome. Based on genome sequencing, 15,009 genes were annotated/identified, compared to the predicted 33,437 genes based on the sequenced transcriptome (predicted by taking into account the number of transcripts less isoforms); this difference in gene number prediction between transcriptomes and genomes is consistent across anaerobic fungi and likely reflective of ploidy.^35,36^ This discrepancy is largely explained by our observation that this strain of *Caecomyces* is likely a diploid (or dikaryon), as we detected ~10k gene models on smaller scaffolds in regions that were >95% identical to regions on larger scaffolds. These scaffolds were designated as secondary scaffolds and these secondary models/alleles were not included in further analyses. Table 1 depicts genomic features for high-resolution sequenced anaerobic fungi, as reported by the JGI MycoCosm pipeline.^37^

**Table 1:**
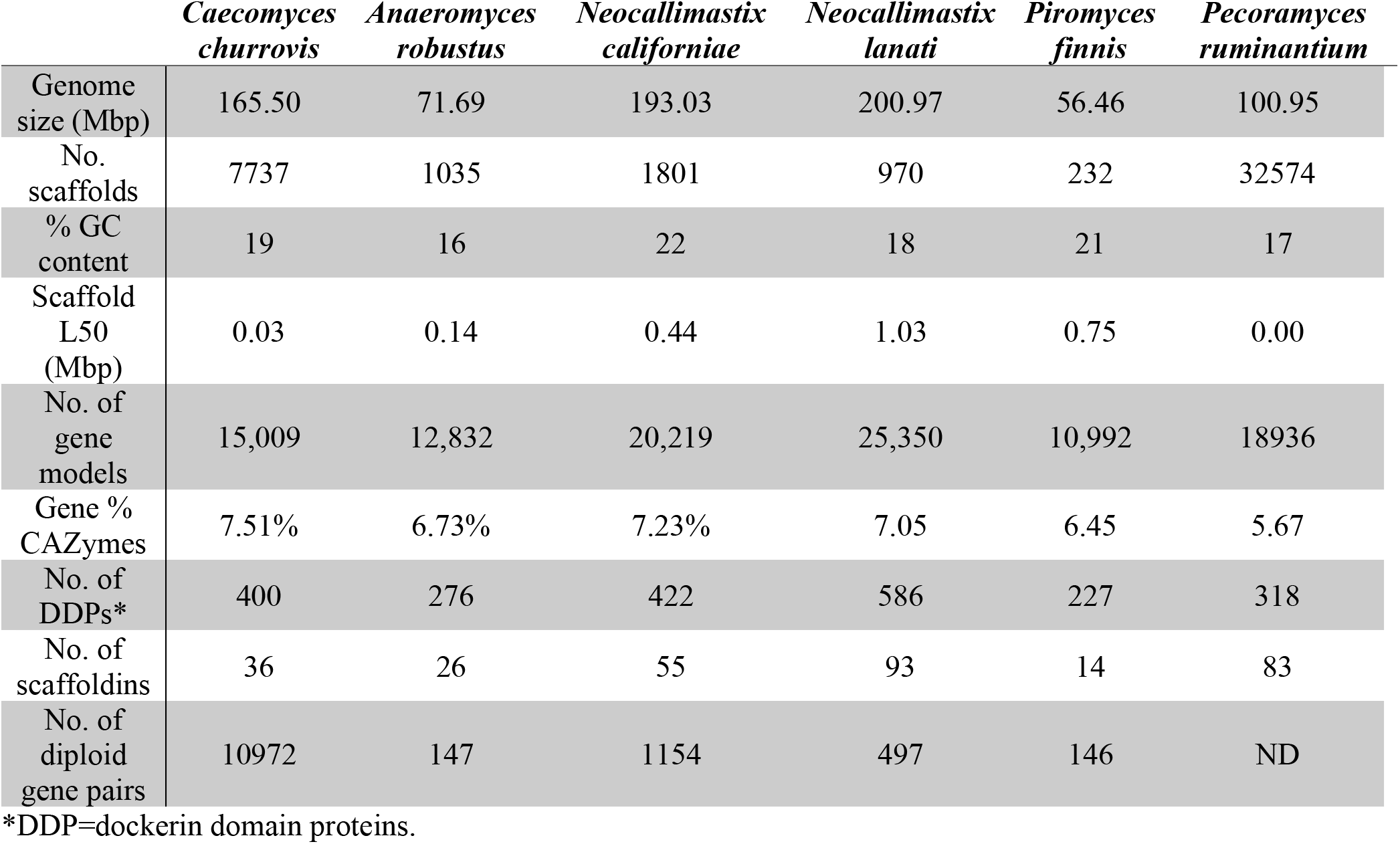
Overview of sequenced anaerobic fungal genome features and statistics^2–4^

As noted in Table 1, the *C. churrovis* genome is GC depleted on the same order of magnitude as the other sequenced anaerobic fungal strains. Such extreme codon biases have made it challenging to heterologously express and evaluate the function of anaerobic fungal genes (like CAZymes) in model systems.^38–40^ Homopolymeric runs of amino acids are found in the *C. churrovis* genomes, which are common in the CAZyme machinery of anaerobic fungi, and could serve as glycosylation sites that prevent proteolytic cleavage.^41^ Collectively, the function of such features need to be better characterized if gut fungal CAZymes from strains such as *C. churrovis* are to be heterologously produced in a model organism.^41^

Anaerobic gut fungi possess an abundance of CAZymes with diverse functions, and are particularly rich in hemicellulases (especially glycosyl hydrolase 10 family) and polysaccharide deacetylases.^32^ Some CAZymes are anchored by non-catalytic fungal dockerin domains (NCDDs) to cohesin domains on large scaffoldin proteins to form enzymatic complexes called fungal cellulosomes.^35^ The high-resolution genome presented here enabled a Hidden Markov Model (HMM) analysis of *C. churrovis* genome, which annotated 36 genes as fungal scaffoldins, compared to the 38 transcripts predicted based on tblastn alignment of the previously sequenced transcriptome. The quantity of predicted proteins identified as cellulases, hemicellulases, and other accessory enzymes along with the total number of CAZymes for each of the 6 sequenced fungal strains are listed in Supplementary Table 1. Fewer total CAZymes in the above categories were identified using predicted proteins found in the sequenced genome (338) than were identified by counting the number of transcripts in the sequenced transcriptome (512), which did not take ploidy into account. The highest abundance accessory enzymes identified in the genome were pectin lyases (15.7% of all CAZymes), in contrast to the transcriptome, in which carbohydrate esterases containing SGNH (defined by four invariant residues – serine, glycine, asparagine, and histidine) hydrolase domains were identified as the most abundant (Supplementary Table 1).^42,43^ However, the *C. churrovis* genome also contains the smallest number of polysaccharide lyase domains (PLs) of any of the 6 fungal genomes characterized (Fig. 1 and Supplementary Table 1).

Proteins containing non-catalytic fungal dockerin domains (NCDDs) were also identified and found to be relatively consistent across strains, in agreement with what was observed for transcript counts (Table 1). However, in contrast to the observation that *C. churrovis* NCDD containing transcripts represented only 15% of all CAZyme transcripts in comparison to 27.9-31.4% for the three other fungal strains examined, the number of NCDD containing proteins represented 30.8% of all CAZyme proteins for *C. churrovis*, similar to the other three fungal strains (Table 1). This suggests that while *C. churrovis* may place greater emphasis on secreted un-complexed, free enzymes to attack plant biomass and release fermentable sugars compared to rhizoid- forming anaerobic fungi based on previously collected transcriptional data, its genome still contains a proportion of NCDD containing proteins similar to that observed in the genomes of rhizoid-forming anaerobic fungal genera. *C. churrovis* also has the second highest number of carbohydrate binding module domains (CBMs) compared to five other high-quality anaerobic fungal genomes (Figure 1). Further analysis revealed that of these genes, *C. churrovis* also possessed the highest number of CBM family 18 domains among anaerobic fungi sequenced to date (Supplementary Fig. 1).

It was previously reported that N6-methyldeoxyadenine (6mA) is associated with transcriptionally active genes in early-diverging fungal lineages in a study using single-molecule long-read sequencing to determine which adenines were methylated.^26^ Of the 6,692 genes that were methylated when the *C. churrovis* genome was sequenced, 4,063 had KOG annotations, 1,002 had KEGG annotations, 3,450 had GO annotations, and 407 were annotated as CAZymes. Almost 1% of all adenines are methylated, and 93% of modifications are at AT dinucleotides, as shown in Supplementary Figure 2A. Very few symmetric runs were present, consistent with avoidance of TAT/ATA reported previously.^26^ Modifications are primarily at the start of genes, specifically ramping up in presence at the start of transcription (Supplementary Figure 2B). 6mA was rare in repetitive regions of the genome (Supplementary Figure 2C) and a large proportion of total 6mA was restricted to genic space (Supplementary Figure 2D).

**Figure 2.**
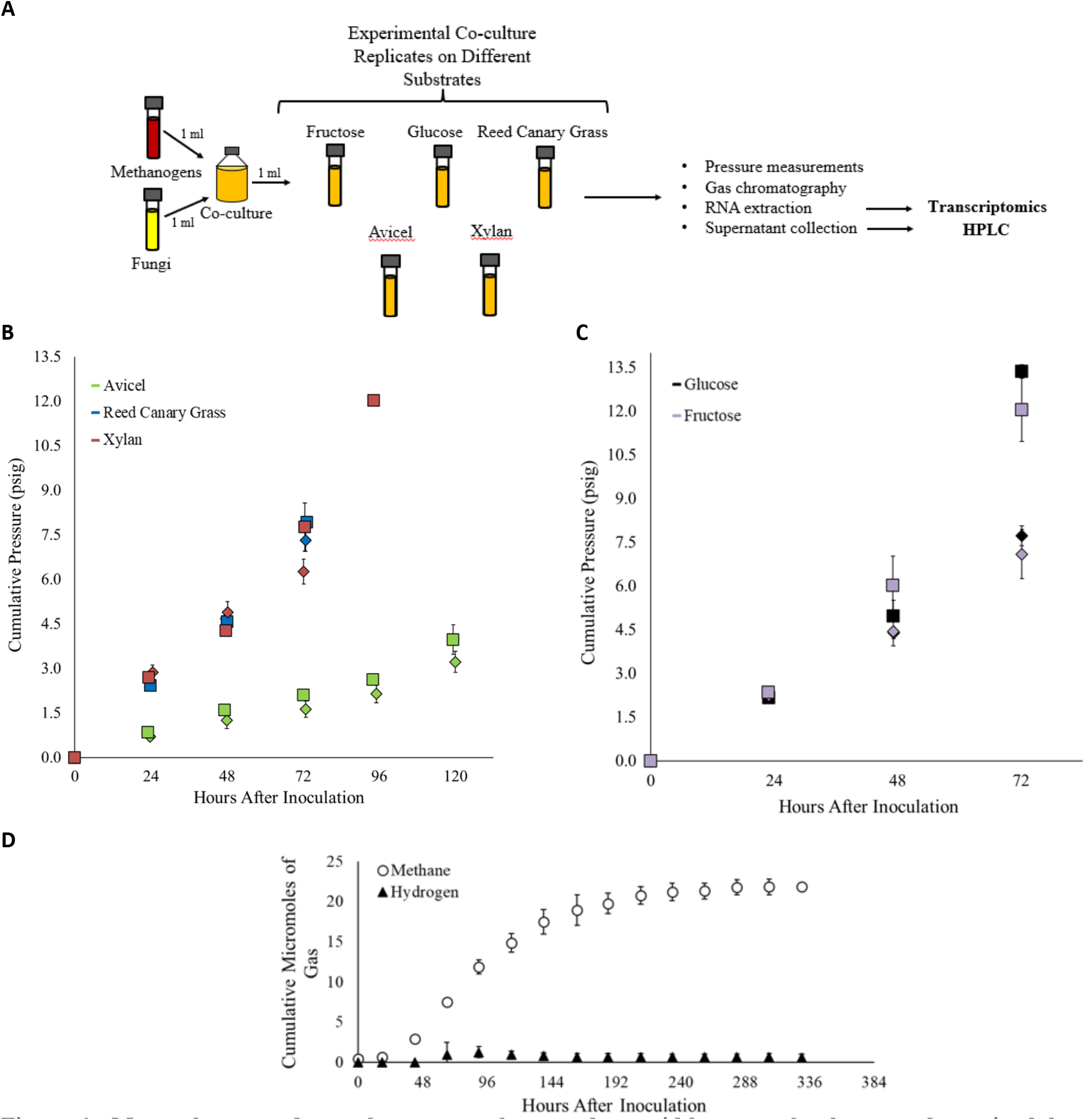
Monocultures and co-cultures were harvested at mid-log growth phase as determined by cumulative pressure. (A) Schematic of the experimental process of cultivating and harvesting co-cultures. A similar process was followed for cultivating and harvesting monocultures, except seed culture was inoculated with 1ml of fungus only.(B and C) Cultures were harvested at pre-determined pressure ranges indicative of the mid-log growth stage for each culturing condition. Cumulative pressure (psig) is plotted versus hours after inoculation for co-cultures and monocultures grown on biomass and components of biomass - reed canary grass, Avicel, and Xylan – in Figure B. Cumulative pressure (psig) is plotted versus hours after inoculation for co-cultures and monocultures grown on soluble sugars – glucose and fructose – in Figure C. Pressure readings for co-cultures are indicated by squares and pressure readings for monocultures are indicated by diamonds. Each substrate is color coded according to the key on the plot. Cultures were harvested at the mid-log growth phase, as indicated by the final pressure time point for each sample. (D) Longterm methane and hydrogen data produced by co-cultures of the anaerobic fungus *C. churrovis* and the methanogen *M. bryantii* on a reed canary grass substrate. Cultures were grown in a complex media formulation, in contrast to cultures harvested for RNA extraction which were grown on MC-. Low levels of accumulated hydrogen indicates stable co-culture over the course of fungal growth.

These results agree with the trends observed for other anaerobic fungal species, further serving to identify 6mA as a widespread epigenetic mark in early-diverging fungi that is associated with transcriptionally active genes.^26^ Note that only ~6% of methylated genes in the genome are annotated as CAZymes, indicating that these genes are not always highly transcribed, but rather the majority of CAZymes are transcribed as needed in response to external stimuli, such as co-culture, growth substrate, etc. Nevertheless, association of gene expression with adenine methylation is necessary to understand and develop transformation techniques, which has proven difficult in anaerobic fungi and other non-model eukaryotic systems to date.^44,45^ Accounting for methylated adenine cluster (MAC) positioning and other epigenetic features could help achieve the methylation required to sufficiently overexpress target genes, such as the CAZymes involved in applications requiring biomass breakdown in both fungal monoculture and in anaerobic biomass-degrading consortia.^26^

### Synthetic co-cultures of C. churrovis with methanogen M. bryantii produce methane

Establishing synthetic co-cultures of anaerobic fungi with methanogens is a valuable tool to probe the impact of co-culture on plant biomass breakdown, substrate uptake, and growth of the individual microbes.^8^ Once plant biomass has been broken down into its constituent sugars by fungal CAZymes, they are catabolized by the fungi and other organisms in the native rumen environment.^46^ Sugars consumed by the fungi undergo glycolysis in the fungal cytoplasm, and the resulting malate and pyruvate are taken up by the fungal hydrogenosome, where they are converted to H_2_ and formate via hydrogenase and pyruvate formate lyase, respectively.^2,47,48^ The hydrogen and formate produced are then exported and available to neighboring methanogens, which assimilate these products and ultimately generate methane.^33^ As such, the metabolic exchange between anaerobic fungi and methanogens benefits both microbes, since it is hypothesized that fungal metabolic end products such as H_2_ and formate may inhibit fungal growth and function if allowed to accumulate, while the methanogens are provided with their required growth substrates.^49^

Figure 2A summarizes the design of this experiment. Cumulative pressure was measured daily (as a proxy for microbial growth) in order to determine when mid-log growth phase had been reached, at which time the cultures were harvested for RNA extraction as shown in Figure 2B and C.^8^ Gas chromatography was used to determine the concentration of methane and hydrogen in the headspace gas of synthetic co-cultures and fungal monocultures on each substrate prior to harvest for RNA extraction at mid-log growth phase. No significant amount of hydrogen was detected in the co-cultures, and no methane was detected in the fungal monocultures, in agreement with *M. bryantii*’s H_2_/CO_2_ requirement for methane production^50^, as shown in Supplementary Figure 3. The absence of hydrogen in the co-cultures indicates that stable pairings of the fungus and methanogen were formed on all substrates (Fig. 2D), which is consistent with previous observations for the *N. californiae* and *A. robustus* anaerobic fungal strains paired with the same methanogen and grown on cellulose and lignocellulosic reed canary grass.^8,9^ Subsequently, transcriptional regulation coupled with HPLC analysis was used to determine the impact of co-cultivation on fungal sugar utilization, hydrogenosome function, secondary metabolite production, and membrane protein regulation in stable, non-native fungal-methanogen co-cultures.

**Figure 3.**
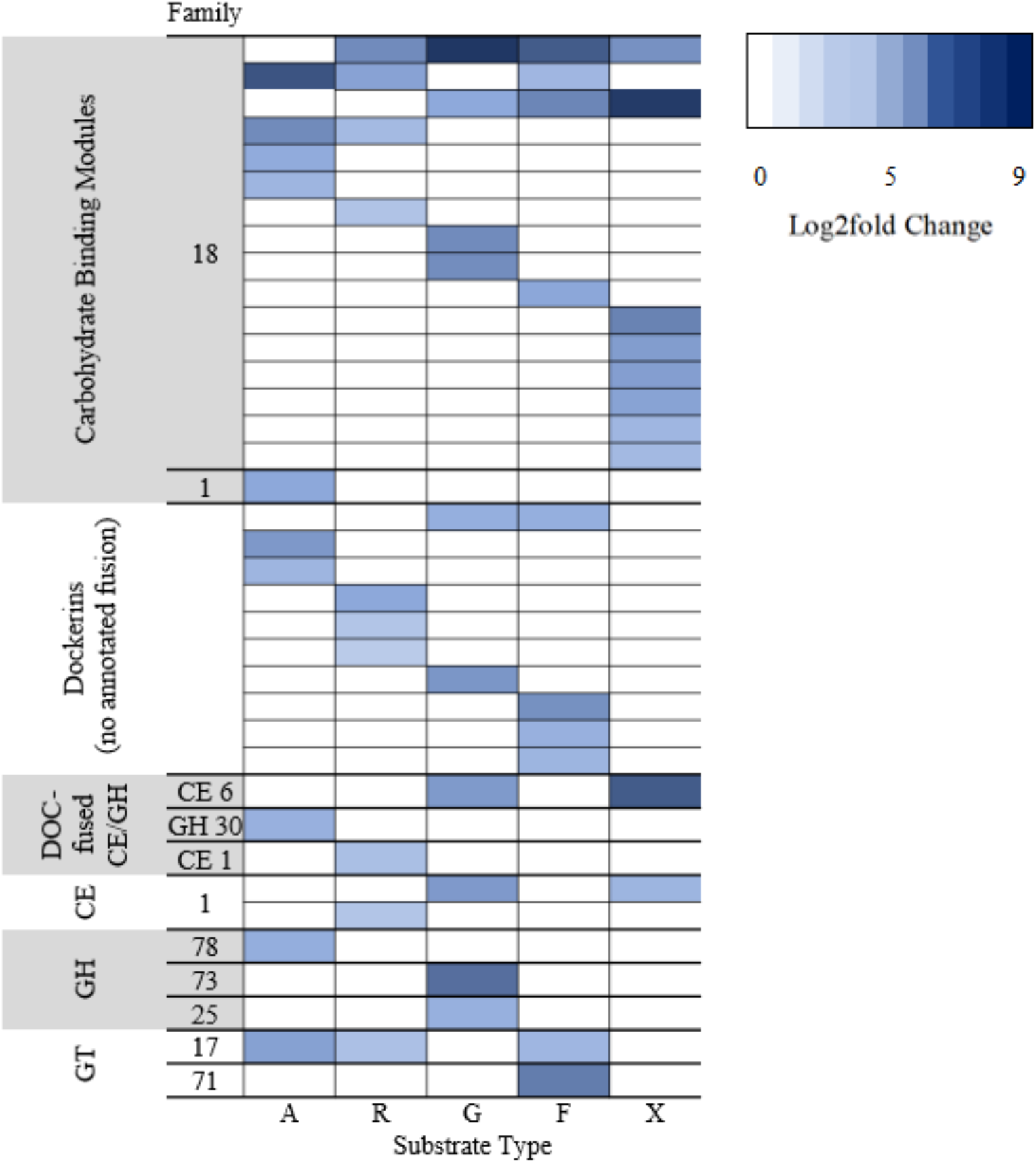
Plot of the top ten upregulated fungal genes annotated as CAZymes or associated enzymatic machinery in co-cultures of the anaerobic fungus *C. churrovis* and the methanogen *M. bryantii* relative to fungal monocultures of *C. churrovis* grown on multiple substrates. Co-cultures of the anaerobic fungus and the methanogen and fungal monocultures were grown on Avicel (A), Reed Canary Grass (R), glucose (G), fructose (F), and Xylan (X). Differential expression of fungal genes in co-cultures relative to fungal monocultures was determined using DESEQ2. The ten genes with the highest log2fold change in expression in co-culture relative to fungal monoculture are shown in the plot above for each substrate and organized into the following classifications: carbohydrate binding modules, dockerins, dockerin-fused carbohydrate esterase or glycoside hydrolases (DOC-fused CE/GH), carbohydrate esterases (CE), glycoside hydrolases (GH), and glycosyltransferases. Protein IDs are listed for each gene. CBMs were highly upregulated, indicating that there may be an increase in enzymatic machinery that aids in anchoring CAZymes to substrates in co-culture, even when grown on soluble sugars. A table containing a list of these genes and the associated log2fold change can be found in Supplementary Table 2.

### Co-culture with a methanogen enhances production of fungal Carbohydrate Binding Modules and fungal dockerins across multiple substrates

Changes in the transcriptional regulation of anaerobic fungi when challenged by different substrates indicates how the fungal CAZyme repertoire and fungal metabolism are adjusted in response to an altered environment. Often, waste streams containing biomass in industrial settings can vary in composition, potentially affecting bioreactor function through shifts in community composition and metabolic function.^51,52^ Examining these changes using RNA-seq reveals how variations in the composition of undefined growth substrates impacts biomass breakdown and product generation. Differential regulation of CAZymes and associated enzymatic machinery was examined for *C. churrovis* co-cultivated with *M. bryantii* and was compared to *C. churrovis* fungal monocultures, both grown on Avicel^®^, reed canary grass, glucose, fructose, and xylan. A proportionally greater number of genes annotated as CAZymes and enzymatic machinery was upregulated in fungal-methanogen co-cultures relative to fungal monocultures than were downregulated on lignocellulose- and hemicellulose-rich substrates, reed canary grass and Avicel^®^. The opposite was true for co-cultures grown on substrates rich in soluble sugars, glucose, fructose and xylan as shown in the Supplementary Figure 4A. The total number of genes upregulated or downregulated for individual CBM, GH, CE, and GT families are shown in Supplemental Figure 4B-E.

**Figure 4.**
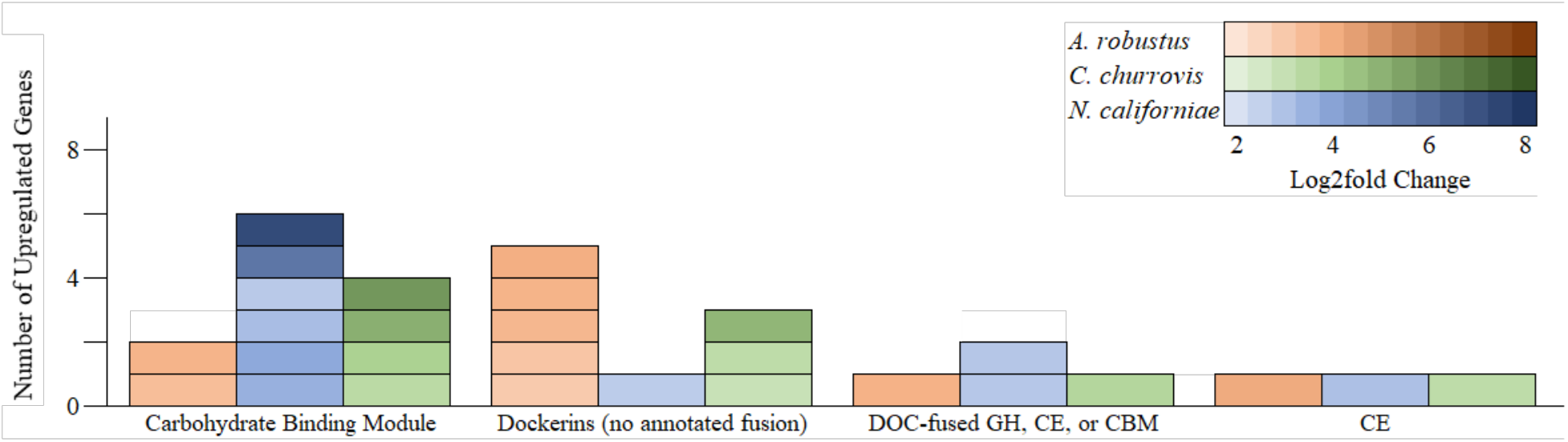
Plot of the top upregulated fungal genes annotated as CAZymes or associated enzymatic machinery in co-cultures of three different fungal strains paired with the same non-native methanogen, *Methanobacterium bryantii* relative to fungal monocultures grown on a reed canary grass substrate. Three different strains of anaerobic fungi, *Anaeromyces robustus*, *Neocallimastix californiae*, and *Caecomyces churrovis* were used to form separate co-cultures with *M. bryantii* and grown on a reed canary grass substrate along with monocultures of each fungus on the same substrate. Differential expression of fungal genes in co-cultures relative to fungal monocultures was determined using DESEQ2. The ten genes with the highest log2fold change in expression in co-culture relative to fungal monoculture are shown in the plot above, with the exception of genes that were not in a category with upregulated genes shared between all three strains (which included one upregulated glycoside hydrolase gene for *A. robustus* and one upregulated glycosyltransferase gene for *C. churrovis*). Genes were organized into the following classifications: carbohydrate binding modules (CBM), DOC (dockerins), dockerin-fused carbohydrate esterase, glycoside hydrolases or carbohydrate binding modules (DOC-fused GH/CE/CBM), and carbohydrate esterases (CE). A table containing a list of these genes and the associated log2fold change can be found in Supplemental Table 3.

However, the majority of the number of top ten genes in these categories upregulated in fungal-methanogen co-culture relative to fungal monoculture on all substrates were annotated as either CBM 18 family proteins or fungal dockerin domains, the majority of which were associated with genes of unknown function. Figure 3 shows the top ten upregulated fungal genes annotated as CAZymes or associated enzymatic machinery in co-cultures of the anaerobic fungus *C. churrovis* and the methanogen *M. bryantii* relative to monocultures of *C. churrovis* grown on multiple substrates. The CBM family with the most abundant number of genes in the sequenced genome, CBM 18, was consistently the gene classification with the greatest log2fold change of any CAZyme orenzymatic machinery on all substrates in fungal-methanogen co-cultures relative to fungal monocultures. Furthermore, the same CBM 18 gene (*Caecomyces churrovis* protein ID 407913) had the greatest log2fold change in fungal-methanogen co-cultures relative to fungal monocultures on reed canary grass, glucose, and fructose substrates. CBM family 18 modules contain approximately 40 amino acid residues and include members with functions linked to modules with chitinase activity or which are lectins.^53,54^ The modules may therefore either be attached to chitinase catalytic domains or in non-catalytic proteins in isolation or as multiple repeats. These carbohydrate-binding proteins possess diversity in ligand specificity and the ability to maintain enzymes in proximity of the substrate, increasing enzyme concentration and potentially leading to more rapid degradation of polysaccharides. These features make these proteins excellent candidates for use in biotechnological applications designed for biomass breakdown.^55,56^

The observation that CAZymes, fungal dockerins, and other biomass degrading machinery are upregulated in all co-cultures, even those grown on glucose is in agreement with previous studies conducted for fungal-methanogen co-cultures on reed canary grass and glucose at mid-log growth stage.^9,10^ Since the majority of the top ten genes upregulated on all substrates were annotated as either CBM 18 family proteins or fungal dockerin domains, this strongly suggests that co-culture with the methanogen *M. bryantii* results in the transcriptional upregulation of enzymatic machinery associated with biomass degradation. Although no transcriptional upregulation of scaffoldin-encoding genes was initially detected in this study, likely due to the more stringent log2fold change cutoff used to determine significant upregulation, Gene Set Enrichment Analysis (GSEA) of the entire set of upregulated genes revealed that upregulated scaffoldins are significantly enriched in co-cultures grown on Avicel^®^ and reed canary grass.^57,58^ These results agree with the finding by Swift, et. al that transcription of fungal cellulosome components increases in co-culture.^59^ Another possibility is that the production of CBM18 transcripts is not related to plant biomass breakdown but instead to interactions between the fungus and methanogen since differential expression is observed across all conditions, including growth on glucose. Many of the dockerin domains not attached to CAZymes contain a CotH kinase protein domain. Previous work showed that approximately 20% of DDPs identified in five previously sequenced anaerobic fungi belonged to spore coat protein CotH and were also present in bacterial cellulosomes.^2^ These dockerin domain proteins belonging to spore coat protein CotH have been speculated to be involved in plant cell wall binding, although this remains to be experimentally validated.^60^

The top ten most highly upregulated genes according to log2fold change annotated as CAZymes, CBMs, or fungal dockerins in co-cultures of *C. churrovis* with *M. bryantii* grown on reed canary grass were compared to those upregulated in co-cultures of the same methanogen, *M. bryantii*, with fungal strains *A. robustus* (previously published) and *N. californiae,* grown on the same substrate.^9^ Of these genes, those falling in categories common to all three strains, which included genes annotated as CBMs, dockerins, and dockerin-fused CAZymes are included in Figure 4. The number of genes regulated in CBM, GT, PL, CE, and GH families in the three fungal strains in co-culture versus fungal monoculture on reed canary grass substrate are shown in Supplementary Figure 5. The most highly upregulated gene for each strain was a CBM family 18 protein for both the *N. californiae* strain and the *C. churrovis* strain and a Carbohydrate Esterase (family 1) protein for the *A. robustus* strain. For each strain, at least three of the top ten genes were fungal dockerin domains, fused to CAZymes or genes of other function. This comparison suggests that co-cultivation with a methanogen likely encourages substrate channeling between synergistic enzymes for both rhizoid-forming fungal strains (*A. robustus* and *N. californiae*) and non-rhizoid-forming fungi (*C. churrovis*).^9,35^ Previously, it was suggested that a smaller proportion of CAZyme transcripts containing dockerin domains in the transcriptome of *C. churrovis* indicated a greater dependence on free enzymes compared to rhizoid-forming gut fungal genera.^5^ Nevertheless, with comparative transcriptomic data, upregulation of these non-catalytic modules and CBMs are clearly observed when *C. churrovis* is cultured with *M. bryantii*. This could indicate that anaerobic fungi, regardless of their usual mode of biomass deconstruction, will respond to the presence of other microbes by increasing binding to fibrous substrates. This would allow them more direct access to sugars released during biomass breakdown, which might otherwise be consumed by other microbes.

**Figure 5.**
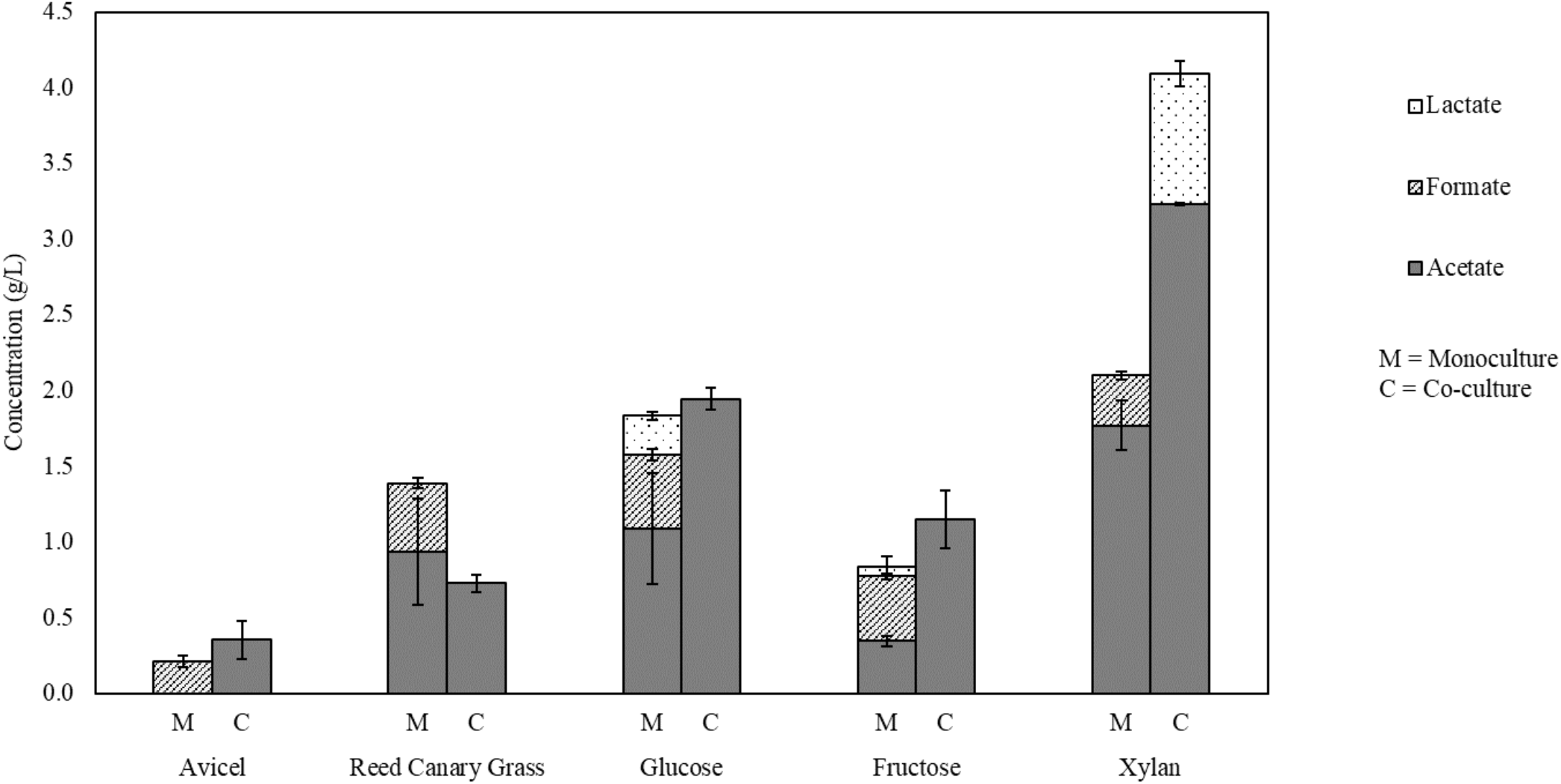
Accumulated metabolites for co-cultures of *C. churrovis* paired with *M. bryantii* versus monocultures of *C. churrovis* upon harvest. HPLC data is shown for co-culture and monoculture grown on each substrate. No formate was observed in co-culture on any substrate, suggesting that *M. bryantii* is capable of metabolizing formate. Trace amounts of ethanol were present in the cultures but fell below the 0.1 g/L limit of detection. This, in conjunction with increased levels of acetate in co-culture, indicates that some of the PFLs upregulated in co-culture in xylan and fructose may be functioning within the hydrogensome.

### Fungal co-culture with a methanogen may enhance PFL function and production of bottleneck enzymes in sugar pathways

Transcriptional regulation coupled with HPLC analysis was used to determine the impact of methanogen co-cultivation on fungal sugar utilization, genes potentially associated with hydrogenosome function, secondary metabolite production, and membrane protein regulation in stable, non-native fungal-methanogen co-cultures. Previous studies of fungal-methanogen co-cultures described increased sugar utilization in co-culture.^14,61^ As such, we hypothesized that genes encoding enzymes involved in sugar catabolism would be upregulated in *C. churrovis* and *M. bryantii* co-cultures relative to fungal monocultures. While some enzymes within these pathways showed changes for each substrate, no co-culture condition resulted in uniform up or downregulation of all enzymes within a given sugar pathway, as shown in Supplementary Figure 6. The enzymes that were upregulated in fungal-methanogen co-culture relative to fungal monoculture on the same substrate may represent bottlenecks in these catabolic pathways. We suspected that sugar utilization in co-cultures could also be increased through upregulation of sugar transporters in the co-culture condition. We instead observe that in the presence of Avicel^®^ and xylan, *M. bryantii* induces transcriptional upregulation of genes that appear to encode proteins homologous to prokaryotic Substrate Binding Proteins (SBPs), as well as Class C G-Protein Coupled Receptors (GPCRs) as seen in Supplemental Table 2.^62–64^ While the function of these protein domains and receptors remains unknown, we speculate that they may be involved in the increased binding of sugar polymers in the presence of the methanogen; or in establishing physical interactions between the methanogens and fungi.^65^

A previous study showed that anaerobic fungal genomes encode a wide array of biosynthetic enzymes of natural products including secondary metabolites - small, bioactive molecules known to mediate a variety of interactions between microorganisms.^66–69^ The majority of these genes were not significantly differentially expressed between co-culture and monoculture conditions on the various substrates in this study. However, two of these fungal genes were highly upregulated in co-culture (*p*-adjusted <0.01). The first is a non-ribosomal peptide synthetase (NRPS)-like gene (protein Id 604712), which was upregulated eight-fold during growth on fructose and on Avicel^®^. The second, a polyketide synthase (PKS; protein Id 402343) was four-fold upregulated in co-culture compared to monoculture during growth on xylan and reed canary grass, suggesting that some fungal secondary metabolites may mediate the interaction between *C. churrovis* and *M. bryantii*, depending on the specific substrate. Co-culture interaction may be most notable on Avicel^®^ and xylan substrates, as both transporters and secondary metabolite biosynthesis genes were upregulated in co-culture for both of these substrates.

Based on previous studies noting an increase in metabolites produced by the ATP-generating fungal hydrogenosome during co-culture with methanogens, we hypothesized that genes associated with hydrogenosomal function would be upregulated in methanogen co-culture.^70,71^ A list of genes associated with the fungal hydrogenosome of the *C. churrovis* strain was constructed based on homology with known hydrogenosome components, shown in Supplemental Table 3. FASTA sequences from known hydrogenosomal components identified in the fungal strain *Neocallimastix lanati*^4^ were aligned to filtered model proteins of *C. churrovis* using the blastp alignment program in MycoCosm. One or more genes within the *C. churrovis* genome aligned to all listed hydrogenosomal enzymes found in *N. lanati*. Regulation of these genes in co-culture compared to monoculture was examined for each substrate. As shown in Supplemental Table 3, 21 genes were homologous to both pyruvate formate lyases (PFLs) that were identified in the *N. lanati* genome.^4^ This enzyme reversibly converts pyruvate and CoA into acetyl-CoA and formate, which plays a central role in anaerobic glucose fermentation.^72^ It has been shown that this enzyme is functional in hydrogenosomes of the anaerobic fungal species *Piromyces* sp. E2 and *Neocallimastix* sp. L2.^73^ The most notable upregulation of PFLs was observed in cultures grown on xylan and fructose, where 15 of the 21 PFL genes identified by homology were upregulated in co-cultures compared to monocultures grown on xylan and two genes identified by homology were upregulated in co-cultures compared to monocultures grown on fructose as shown in Supplementary Table 3. Five additional genes annotated as PFLs (or formate C acetyltransferases) according to Enzyme Commission (EC) number rather than homology to the *N. lanati* genome were upregulated on xylan and one additional gene was upregulated on fructose. One of these genes (Protein ID 428490) was upregulated in co-culture on all substrates examined except reed canary grass. A previous study examining transcriptional regulation of co-cultures of the native fungus-methanogen pairing *Pecoramyces* sp. F1 with the methanogen *Methanobrevibacter thaueri* versus monoculture of the fungus grown on glucose did not detect a difference in expression levels of PFL genes (although upregulation was detected at the protein level).^10^

Although we hypothesized that genes associated with the hydrogenosome would be transcriptionally upregulated in the co-culture relative to the fungal monocultures based on the metabolic data collected in previous work, transcriptional upregulation of genes associated with hydrogenosomal function is limited, with the exception of pyruvate formate lyases in co-cultures grown on xylan and fructose. It is important to note that further studies are needed to confirm that this transcriptional upregulation of PFLs is associated specifically with the hydrogenosome, as PFLs function in both the cytosol and the hydrogenosome. However, as a complement to the transcriptional information regarding metabolic function in this study, end point metabolites present in the supernatant were measured using HPLC upon harvest of the co-cultures and monocultures (Figure 5). Increases in the amount of acetate produced in co-culture and the absence of significant amounts of ethanol and lactate indicate that some of these genes may potentially be associated with hydrogenosome function for cultures grown on fructose, since pyruvate can either be converted to lactate or ethanol by PFLs functioning in the cytosol or converted to acetate by PFLs functioning within the hydrogenosome. Ethanol was also absent in cultures grown on xylan, although higher levels of lactate were observed in co-culture in addition to higher levels of acetate, indicating that both cytosolic and hydrogenosomal PFLs may be upregulated in co-culture. GSEAPreranked analysis also indicated that upregulated genes were enriched in pathways associated with pyruvate metabolism and glycolysis for co-cultures grown on xylan, in agreement with the observed upregulation of PFLs.^57,58^

While analysis of the end-point metabolites of *A. robustus* paired with *M. bryantii* in previous work did not indicate a statistically significant difference in the level of formate in co-culture versus monoculture, formate was absent in the *C. churrovis* and *M. bryantii* co-culture samples but present in fungal monocultures.^9^ Earlier studies concluded that this type strain of *M. bryantii* (DSM 863 M.o.H.) was unable to produce methane from formate in pure culture.^74,75^ The discovery of a formate transporter and several copies of formate dehydrogenase genes upon sequencing the methanogen’s genome has suggested the possibility of growth on formate.^50^ The observed upregulation of PFL genes and the absence of formate in co-cultures in the current study provides evidence that this strain of *M. bryantii* can utilize formate under certain conditions. A similar phenomenon has been observed for co-cultivation of a formate-producing *Piromyces* fungal species and the natively associated methanogen *Methanobrevibacter thaueri*, a methanogen that has been shown incapable of growth on formate.^20,76^ It is possible that cultivating these methanogens under the conditions required for co-culture with rumen anaerobic fungi stimulate formate utilization by inducing function of the formate transporter and formate dehydrogenase discovered upon sequencing the genome.^50^

## Conclusions

Here, we have sequenced the first high-quality genome of a non-rhizoidal fungus, *Caecomyces churrovis*, revealing an abundance of diverse CAZymes and the highest number of CBM family 18 domains among anaerobic fungi sequenced to date. We found that co-cultivation of the *C. churrovis* fungal strain with the non-native methanogen *M. bryantii* enhanced production of transcripts containing these chitin-binding CBM 18 domains across a variety of substrates. Upregulation of CBMs and dockerin domains in fungal-methanogen co-culture with the same non-native methanogen relative to fungal monoculture on a lignocellulose-rich substrate was also observed for two other previously sequenced fungal strains, *A. robustus* and *N. californiae*. We hypothesize that the function of CBMs belonging to family 18 may not be directly related to plant biomass breakdown but instead to interactions between the fungus and methanogen since upregulation of transcripts containing these domains is observed across multiple cultivation conditions, including both cellulose and lignocellulose-rich substrates as well as soluble sugars. Upregulation of genes associated with sugar pathways and the functioning of the hydrogenosome for *C. churrovis* and *M. bryantii* co-cultures relative to fungal monocultures of *C. churrovis* also suggests that co-culture with a methanogen may enhance pyruvate formate lyase (PFL) function under certain cultivation conditions and production of key enzymes in sugar utilization pathways. Overall, these observations enhance our understanding of the mechanistic interactions between anaerobic fungi and associated methanogens, which aids in our ability to design synthetic biomass-degrading microbial consortia.

## Acknowledgements

We thank the following for funding support: the National Science Foundation (NSF, grant no. MCB-1553721), the Office of Science (BER) of the US Department of Energy (DOE) (grant no. DE-SC0010352), the Institute for Collaborative Biotechnologies (grant nos. W911NF-09–D-0001, W911NF-19-2-0026, and W911NF-19-1-0010) from the US Army Research Office, and the Camille Dreyfus Teacher-Scholar Awards Program. The sequencing conducted by the US DOE Joint Genome Institute, a DOE Office of Science User Facility, is supported by the Office of Science of the US DOE under contract no. DE-AC02-05CH11231. The authors thank Patrick Leggieri, Stephen Lillington, and Amy Eisenberg for helpful discussions and revision of the manuscript.

