## Supplementary Information for "Co‑cultivation of the anaerobic fungus *Caecomyces churrovis* with *Methanobacterium bryantii* enhances transcription of carbohydrate binding modules"

**Supplemental Information**

**Supplementary Table 1.** The number of proteins identified as cellulases, hemicellulases, and other accessory enzymes for six sequenced anaerobic fungi annotated from genome sequencing (see methods).

| | Number of protein IDs | | | | | |
|---|---|---|---|---|---|---|
| **Hemicellulases** | *C. churrovis* | *N. californiae* | *A. robustus* | *P. finnis* | *N. lanati* | *O. sp. C1A* |
| GH11 | 33 | 24 | 29 | 36 | 90 | 45 |
| GH43 | 25 | 43 | 18 | 14 | 48 | 31 |
| GH10 | 12 | 58 | 15 | 21 | 59 | 32 |
| GH39 | 7 | 9 | 5 | 2 | 8 | 3 |
| GH30 | 4 | 4 | 2 | 1 | 4 | 3 |
| **Accessory enzymes** | | | | | | |
| Carbohydrate esterase (SGNH hydrolase domains) | 51 | 68 | 39 | 23 | 79 | 39 |
| Pectin Lyase | 53 | 127 | 44 | 49 | 142 | 72 |
| Polysaccharide deacetylase | 43 | 93 | 49 | 44 | 96 | 48 |
| Rhamnogalacturonate lyase | 2 | 9 | 3 | 2 | 12 | 1 |
| Pectinesterase | 1 | 15 | 5 | 5 | 16 | 8 |
| Glycosyl Hydrolase 88 | 0 | 2 | 0 | 1 | 2 | 0 |
| **Cellulases** | | | | | | |
| GH9 | 10 | 14 | 9 | 12 | 14 | 13 |
| GH6 | 18 | 27 | 12 | 21 | 89 | 49 |
| GH45 | 20 | 28 | 14 | 15 | 28 | 16 |
| GH48 | 7 | 21 | 7 | 13 | 23 | 14 |
| GH1 | 7 | 16 | 7 | 10 | 19 | 10 |
| GH5 | 23 | 65 | 26 | 26 | 70 | 47 |
| GH3 | 10 | 53 | 15 | 15 | 58 | 18 |
| GH16 | 8 | 19 | 11 | 6 | 19 | 6 |
| GH8 | 1 | 2 | 2 | 1 | 2 | 1 |
| GH31 | 3 | 10 | 7 | 2 | 11 | 19 |
| Total | 338 | 706 | 319 | 319 | 889 | 475 |

**Supplemental Table 2**. Co-cultivation of *C. churrovis* with *M. bryantii* induces transcriptional upregulation of genes that appear to encode proteins homologous to prokaryotic Substrate Binding Proteins (SBPs), as well as Class C G-Protein Coupled Receptors (GPCRs). Among regulated transcripts are sequences encoding G-protein coupled receptors (7tm_3, PF00003) and sequences encoding putative Substrate Binding Proteins (SBP_Bac_1, PF01547; SBP_Bac_3, PF00497; and SBP_Bac_8, PF13416). The table lists transcriptional regulation of genes encoding proteins that have at least one predicted transmembrane segment. Approximately half of all sequences contain at least one Pfam. Conditions indicate the substrate on which cultures were grown, followed by whether *C. churrovis* transcripts were upregulated or downregulated in the co-culture condition relative to fungal monocultures.

| Condition | Number of transcripts affected | Transcripts with at least 1 Pfam hit | GPCR | SBP |
|---|---|---|---|---|
| Glucose Upregulated | 149 | 77 | 1 | 0 |
| Glucose Downregulated | 203 | 123 | 5 | 21 |
| Fructose Upregulated | 87 | 48 | 1 | 0 |
| Fructose Downregulated | 66 | 33 | 1 | 5 |
| Avicel Upregulated | 321 | 183 | 21 | 41 |
| Avicel Downregulated | 22 | 15 | 0 | 1 |
| Xylan Upregulated | 175 | 101 | 15 | 8 |
| Xylan Downregulated | 278 | 164 | 1 | 2 |
| Reed Canary Grass Upregulated | 44 | 17 | 0 | 0 |
| Reed Canary Grass Downregulated | 18 | 13 | 0 | 0 |

**Supplementary Table 3.** One or more genes within the *C. churrovis* genome aligned to all listed hydrogenosomal enzymes for *N. lanati*, with a %identity cutoff of 70.0 and a %subject coverage cutoff of 80.0, with the exception of the complex 2 subunit D, which only had a %subject coverage of 59.0 (%identity was 79.6)

| ***C. churrovis*** **Protein ID** | **Enzyme** | **Upregulated or Downregulated (Substrate)** |
|---|---|---|
| 428490 | PFL1/PFL2 | Upregulated (X, A, G, F) |
| 11340 | PFL1/PFL2 | Upregulated (X) Downregulated (A) |
| 193710 | PFL1/PFL2 | Upregulated (X) Downregulated (R) |
| 193705 | PFL1/PFL2 | Upregulated (X) Downregulated (R) |
| 420504 | PFL1/PFL2 | |
| 107174 | PFL1/PFL2 | Upregulated (X, G) |
| 417119 | PFL1/PFL2 | |
| 621094 | PFL1/PFL2 | Upregulated (X) |
| 431187 | PFL1/PFL2 | Upregulated (X) |
| 214975 | PFL1/PFL2 | Upregulated (X, F) |
| 621093 | PFL1/PFL2 | Upregulated (X) |
| 277171 | PFL1/PFL2 | Upregulated (X) |
| 622192 | PFL1/PFL2 | Upregulated (X) Downregulated (G) |
| 277169 | PFL1/PFL2 | Upregulated (X) |
| 418039 | PFL1/PFL2 | Upregulated (X) |
| 416923 | PFL1/PFL2 | Upregulated (X) |
| 537129 | PFL1/PFL2 | Upregulated (X) |
| 635526 | PFL1/PFL2 | |
| 413357 | PFL1/PFL2 | |
| 572185 | PFL1/PFL2 | |
| 462330 | PFL1/PFL2 | |
| 454624 | Ac:SucCoA trans. | |
| 454624 | Ac:SucCoA trans. | |
| 453440 | SucCoA syn. Sub. A | |
| 519176 | SucCoA syn. Sub. B | |
| 526974 | SucCoA syn. Sub. B | |
| 243125 | Hydrogenase 1 | Downregulated (R, G) |
| 456067 | Hydrogenase 2 | |
| 557447 | Complex 1: nuoF | Downregulated (G) |
| 454874 | Complex1: nuoE | Downregulated (X, G) |
| 452336 | Complex 2: sub. A | |
| 416671 | Complex 2: sub. B | |
| 544208 | Complex 2: sub. C | |
| 417861 | Complex 2: sub. D | |
| 549900 | Fumarase | |
| 487810 | ATP syn.: sub. Alpha | |
| 443140 | ATP syn: sub. Beta | |
| 459763 | ATP syn: sub. Delta | |
| 523564 | ATP syn.: sub. Gamma | Upregulated (G) |

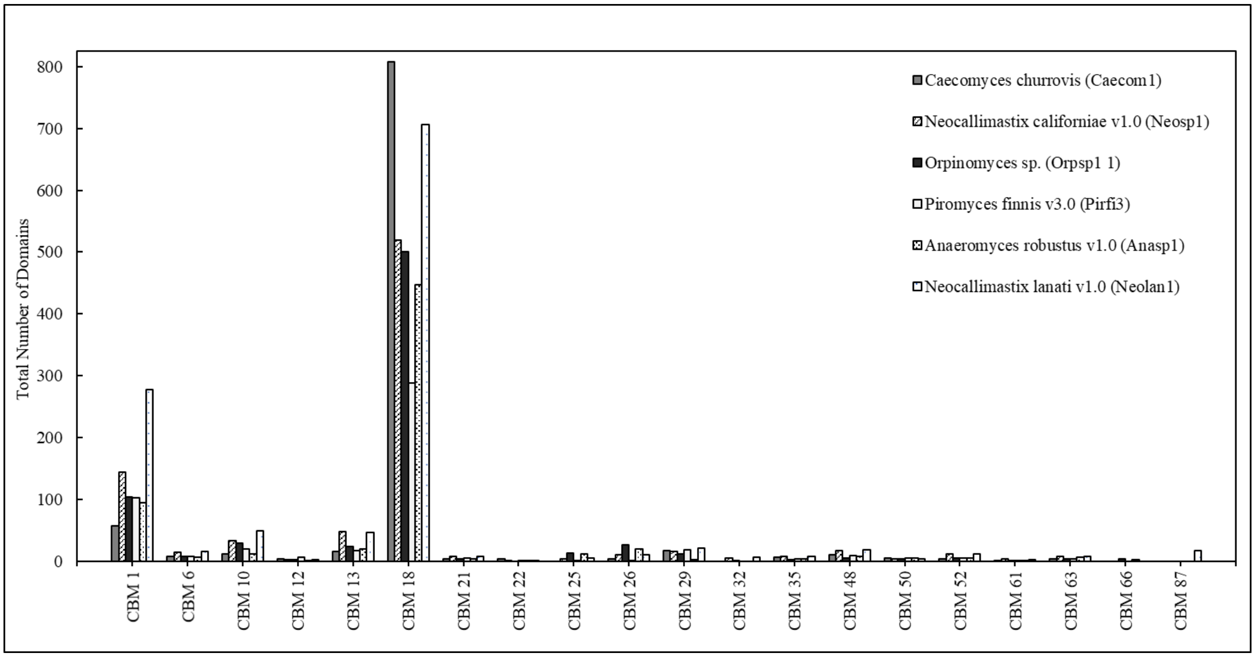


**Supplementary Figure 1.** *Caecomyces churrovis* has the highest number of CBM family 18 domains among sequenced anaerobic fungi to date.

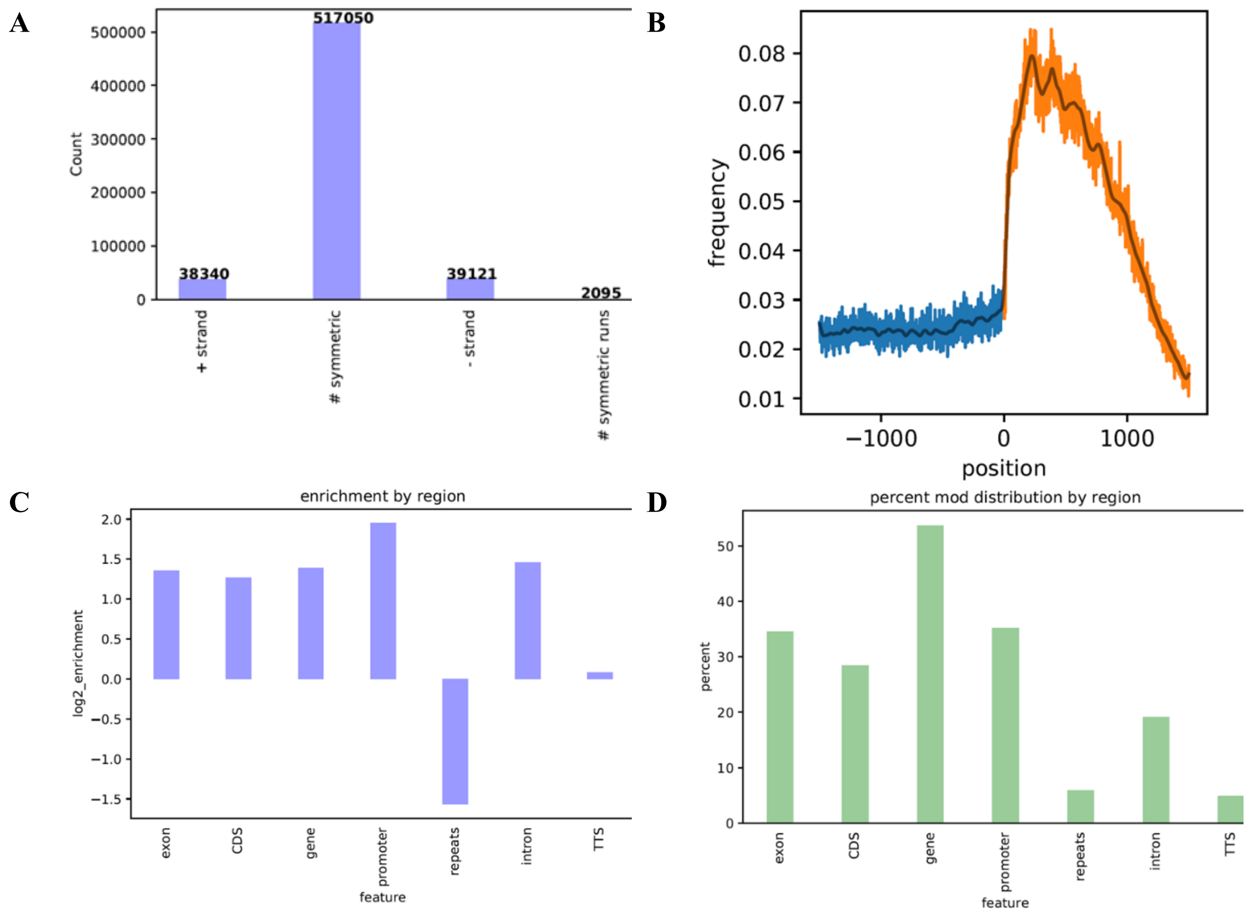


**Supplementary Figure 2. 6mA Modifications occur symmetrically at ApT dinucleotides and are concentrated in methylated adenine clusters (MACs) surrounding the transcriptional start sites of expressed genes.** This finding agrees with previous work examining 6mA modification in 16 other fungal genomes, including other early-diverging fungi. As shown in Figure 2A, 92.2% of modifications were symmetric within AT context and 83.32% of all modifications were symmetric. In addition, 89.67% of modifications were in AT context. Figure 2B shows the frequency (# 6mA observed / # available sites) per position ±1500 bp surrounding transcriptional start sites in *C. churrovis*. A slight wave in modification frequency is observed following the start of the 5' utr, not seen before in other early-diverging fungi, but the presence of modifications at the start of genes is in agreement with previous studies of fungal 6mA modifications.[44] Figure 2C shows the log2fold enrichment of 6mA modifications by region. Log2_enrichment refers to the enrichment of 6mA at a given feature relative to the expected abundance of 6mA genome-wide, normalized by GC content. Figure 2D shows the percent of total 6mA modifications that are found within each region. Note that many of these regions overlap, such as introns and genes. Regions are defined as follows: exons – non-intron genic space, CDS – coding sequence only, gene – entirety of genic space, promoter – ±500bp surrounding transcriptional start sites, repeats – repetitive sequences identified using RepeatScout and RepeatMasker, intron – introns within genic regions, TTS – ±250bp surrounding transcription termination sites.

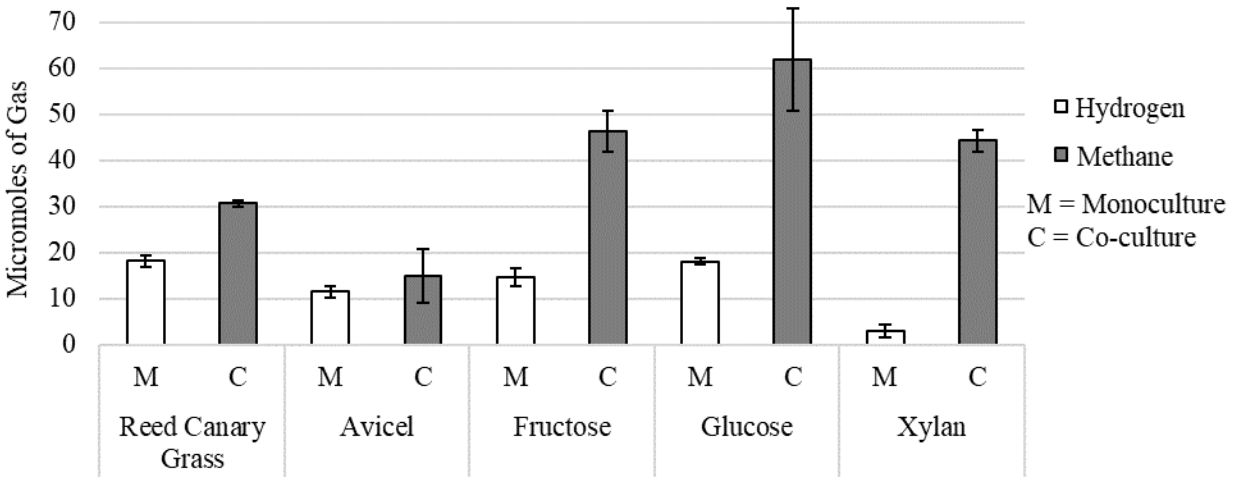


**Supplementary Figure 3.** End-point methane and hydrogen measurements for monocultures and co-cultures. Gas chromatography was used to determine the concentration of methane and hydrogen in the headspace gas of co-cultures and monocultures on each substrate upon harvest for RNA extraction. No significant amount of hydrogen was detected in the co-cultures, and no methane was detected in the monocultures. Significantly higher amounts of methane were produced on soluble substrates.

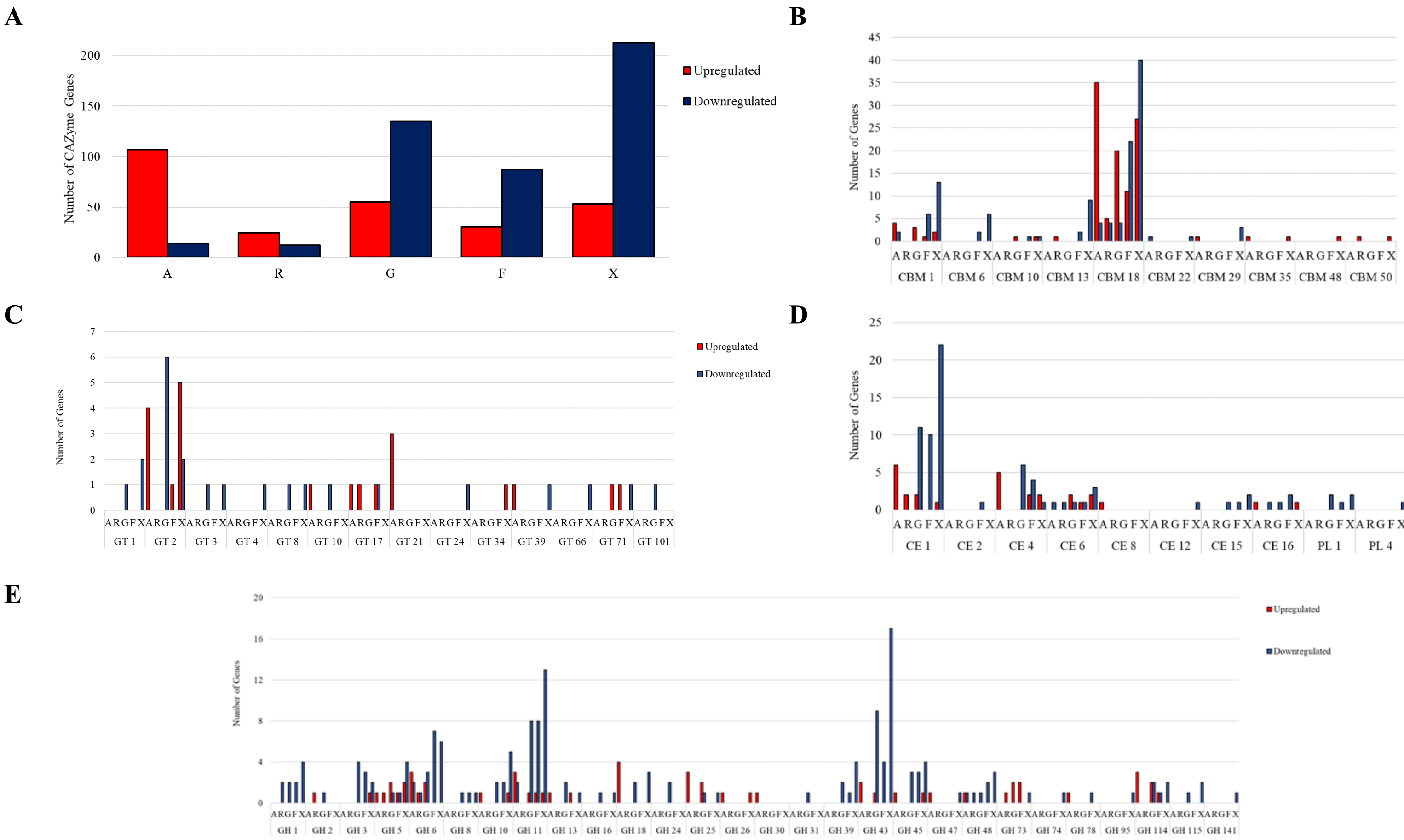


**Supplementary Figure 4.** The total number of genes annotated as CAZymes upregulated or downregulated overall and for CBM, GH, CE, and GT families for fungal-methanogen co-cultures of *C. churrovis* paired with *M. bryantii* relative to fungal monocultures of C. churrovis on a range of substrates. A=Avicel, R=Reed Canary Grass, G=Glucose, F=Fructose, X=Xylan. CBM=Carbohydrate Binding Module, GT=Glycosyltransferase, GH=Glycoside Hydrolase, CE=Carbohydrate Esterase, PL=Polysaccharide Lyase.

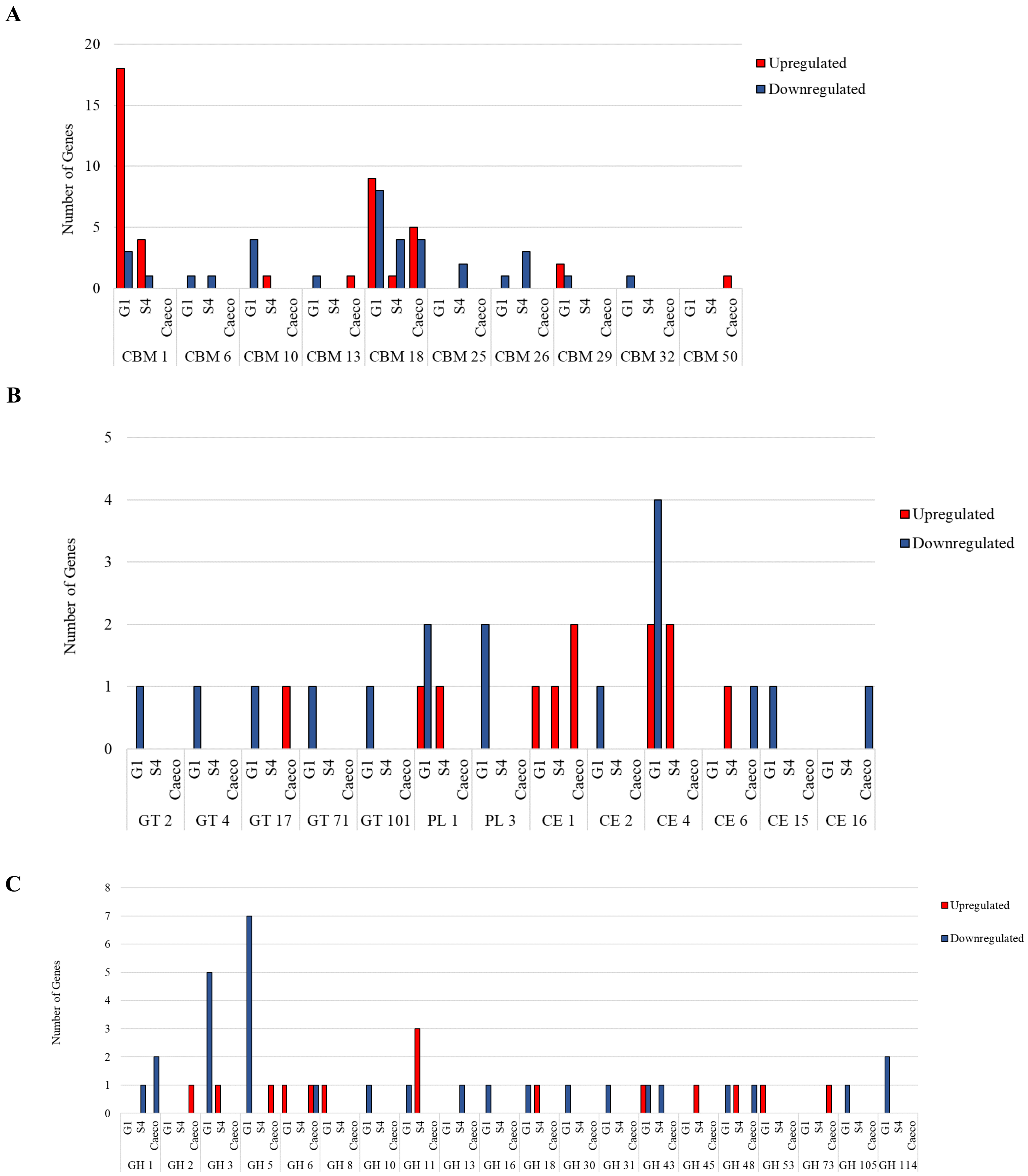


**Supplementary Figure 5. Regulated CBM, GT, PL, CE, and GH families in three fungal strains in co-culture vs monoculture on a Reed Canary Grass Substrate.** CBM 1 and CBM 18 families were significantly regulated (number of genes regulated > =5). The CBM 1 family was highly upregulated for the *N. californiae* strain. GH 3 and GH 5 were significantly downregulated in the *N. californiae* strain (number of genes > or =5). GH=glycoside hydrolase, GT=glycosyltransferase, PL=polysaccharide lyase, CBM=carbohydrate binding module, CE=carbohydrate esterase, Caeco=*Caecomyces churrovis*, S4=*Anaeromyces robustus*, G1=*Neocallimastix californiae*.

**A**

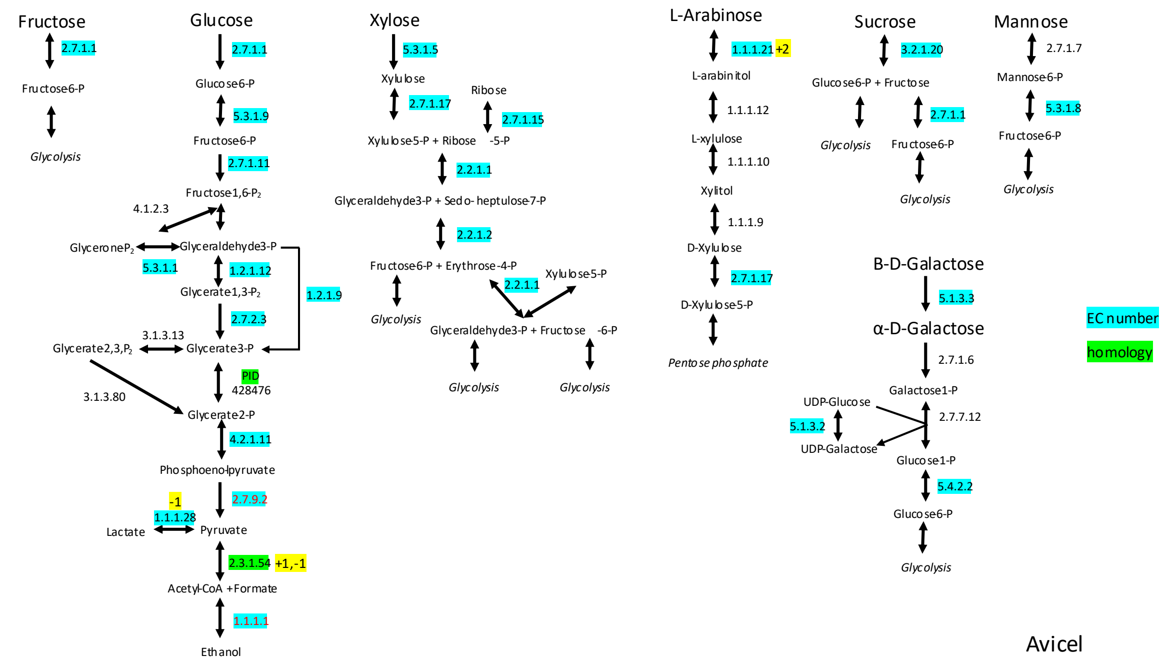
Fructose
2.7.1.1
Fructose6-P
Glycolysis
Glucose
2.7.1.1
Glucose6-P
5.3.1.9
Fructose6-P
2.7.1.11
Fructose1,6-P$_2$
4.1.2.3
GlyceroneP$_2$
Glyceraldehyde3-P
5.3.1.1
1.2.1.12
1.2.1.9
Glycerate1,3-P$_2$
2.7.2.3
3.1.3.13
Glycerate2,3,P$_2$
Glycerate3-P
PID
428476
3.1.3.80
Glycerate2-P
4.2.1.11
Phosphoenolpyruvate
-1
2.7.9.2
1.1.1.28
Lactate
Pyruvate
2.3.1.54
+1,-1
Acetyl-CoA +Formate
1.1.1.1
Ethanol
Xylose
5.3.1.5
Xylulose
2.7.1.17
Ribose
2.7.1.15
Xylulose5-P + Ribose -5-P
2.2.1.1
Glyceraldehyde3-P + Sedo- heptulose7-P
2.2.1.2
Fructose6-P + Erythrose-4-P
Xylulose5-P
2.2.1.1
Glycolysis
Glyceraldehyde3-P + Fructose -6-P
Glycolysis
Glycolysis
L-Arabinose
1.1.1.21
+2
L-arabinitol
1.1.1.12
L-xylulose
1.1.1.10
Xylitol
1.1.1.9
D-Xylulose
2.7.1.17
D-Xylulose5-P
Pentose phosphate
Sucrose
3.2.1.20
Glucose6-P + Fructose
2.7.1.1
Glycolysis
Fructose6-P
Glycolysis
Mannose
2.7.1.7
Mannose6-P
5.3.1.8
Fructose6-P
Glycolysis
B-D-Galactose
5.1.3.3
α-D-Galactose
2.7.1.6
Galactose1-P
UDP-Glucose
2.7.7.12
5.1.3.2
UDP-Galactose
Glucose1-P
5.4.2.2
Glucose6-P
Glycolysis
EC number
homology
Avicel

**B**

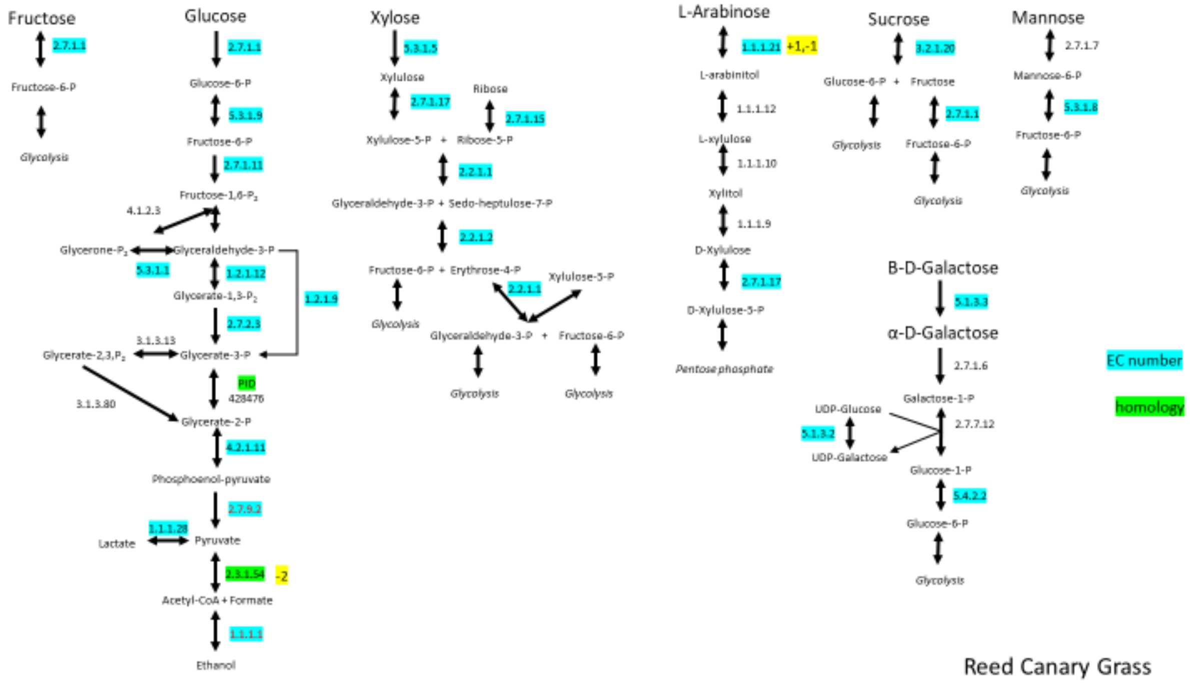
Fructose
Glucose
Xylose
L-Arabinose
+1,-1
Sucrose
Mannose
B-D-Galactose
α-D-Galactose
EC number
homology
-2
Reed Canary Grass

**C**

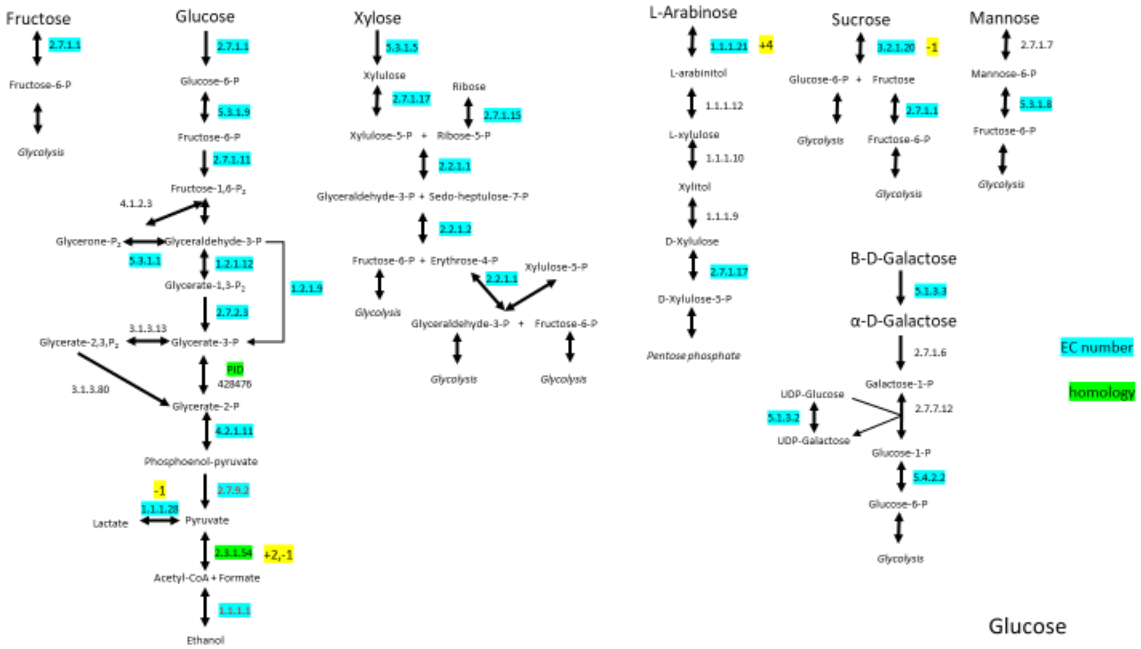

Fructose
2.7.1.1
Fructose-6-P
Glycolysis
Glucose
2.7.1.1
Glucose-6-P
5.3.1.9
Fructose-6-P
2.7.1.11
Fructose-1,6-P2
4.1.2.3
Glycerone-P2
Glyceraldehyde-3-P
5.3.1.1
1.2.1.12
1.2.1.9
Glycerate-1,3-P2
2.7.2.3
3.1.3.13
Glycerate-2,3,P2
Glycerate-3-P
PID
428476
3.1.3.80
Glycerate-2-P
4.2.1.11
Phosphoenol-pyruvate
-1
2.7.9.2
1.1.1.28
Lactate
Pyruvate
2.3.1.54
+2,-1
Acetyl-CoA + Formate
1.1.1.1
Ethanol
Xylose
5.3.1.5
Xylulose
2.7.1.17
Ribose
2.7.1.15
Xylulose-5-P + Ribose-5-P
2.2.1.1
Glyceraldehyde-3-P + Sedo-heptulose-7-P
2.2.1.2
Fructose-6-P + Erythrose-4-P
2.2.1.1
Xylulose-5-P
Glycolysis
Glyceraldehyde-3-P + Fructose-6-P
Glycolysis
Glycolysis
L-Arabinose
1.1.1.21
+4
L-arabinitol
1.1.1.12
L-xylulose
1.1.1.10
Xylitol
1.1.1.9
D-Xylulose
2.7.1.17
D-Xylulose-5-P
Pentose phosphate
Sucrose
3.2.1.20
-1
Glucose-6-P + Fructose
2.7.1.1
Glycolysis
Fructose-6-P
Glycolysis
Mannose
2.7.1.7
Mannose-6-P
5.3.1.8
Fructose-6-P
Glycolysis
B-D-Galactose
5.1.3.3
α-D-Galactose
2.7.1.6
Galactose-1-P
UDP-Glucose
2.7.7.12
5.1.3.2
UDP-Galactose
Glucose-1-P
5.4.2.2
Glucose-6-P
Glycolysis
EC number
homology
Glucose


**D**

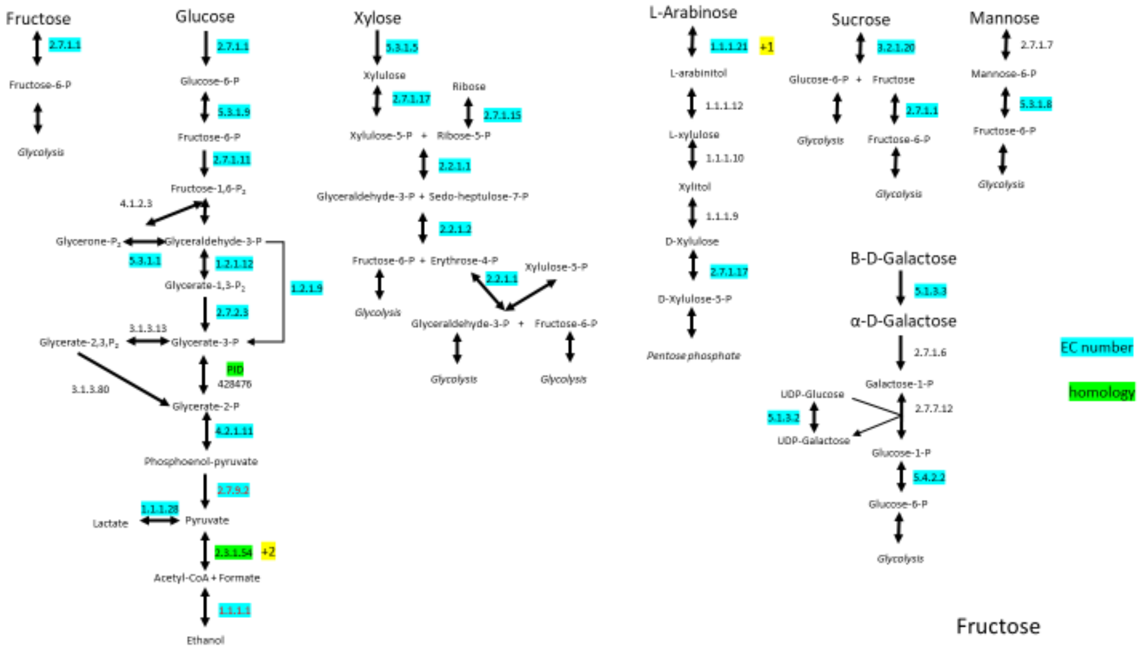

Fructose
2.7.1.1
Fructose-6-P
Glycolysis
Glucose
2.7.1.1
Glucose-6-P
5.3.1.9
Fructose-6-P
2.7.1.11
Fructose-1,6-P2
4.1.2.3
Glycerone-P2
Glyceraldehyde-3-P
5.3.1.1
1.2.1.12
1.2.1.9
Glycerate-1,3-P2
2.7.2.3
3.1.3.13
Glycerate-2,3,P2
Glycerate-3-P
PID
428476
3.1.3.80
Glycerate-2-P
4.2.1.11
Phosphoenol-pyruvate
2.7.9.2
1.1.1.28
Lactate
Pyruvate
2.3.1.54
+2
Acetyl-CoA + Formate
1.1.1.1
Ethanol
Xylose
5.3.1.5
Xylulose
2.7.1.17
Ribose
2.7.1.15
Xylulose-5-P + Ribose-5-P
2.2.1.1
Glyceraldehyde-3-P + Sedo-heptulose-7-P
2.2.1.2
Fructose-6-P + Erythrose-4-P
2.2.1.1
Xylulose-5-P
Glycolysis
Glyceraldehyde-3-P + Fructose-6-P
Glycolysis
Glycolysis
L-Arabinose
1.1.1.21
+1
L-arabinitol
1.1.1.12
L-xylulose
1.1.1.10
Xylitol
1.1.1.9
D-Xylulose
2.7.1.17
D-Xylulose-5-P
Pentose phosphate
Sucrose
3.2.1.20
Glucose-6-P + Fructose
2.7.1.1
Glycolysis
Fructose-6-P
Glycolysis
Mannose
2.7.1.7
Mannose-6-P
5.3.1.8
Fructose-6-P
Glycolysis
B-D-Galactose
5.1.3.3
α-D-Galactose
2.7.1.6
Galactose-1-P
UDP-Glucose
2.7.7.12
5.1.3.2
UDP-Galactose
Glucose-1-P
5.4.2.2
Glucose-6-P
Glycolysis
EC number
homology
Fructose

**E**

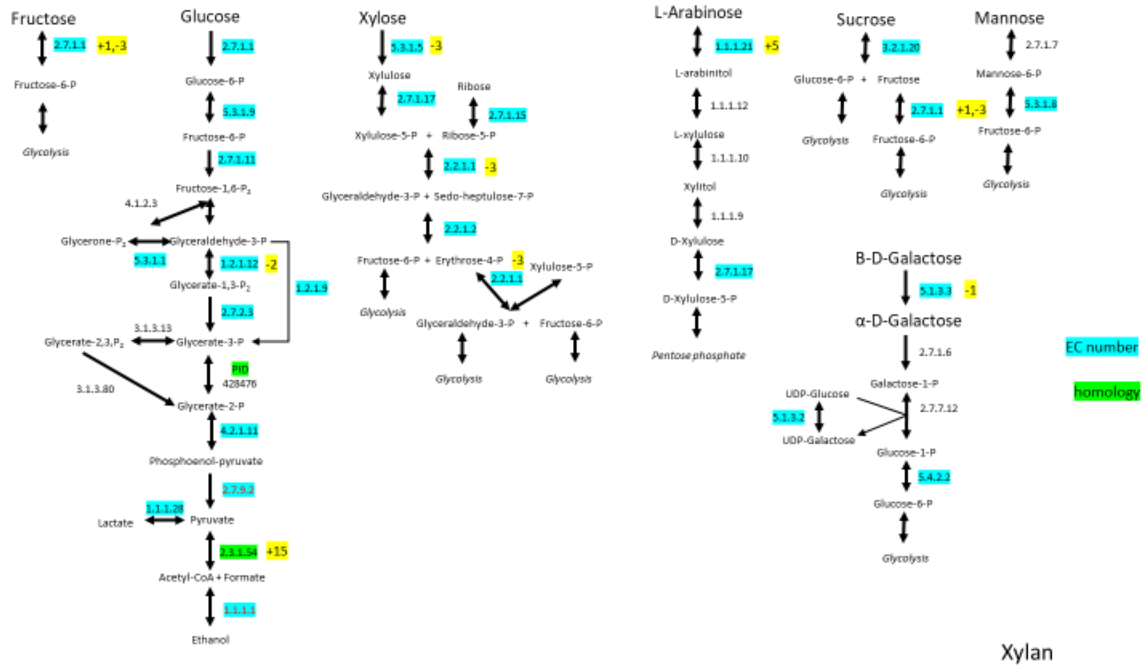


**Supplementary Figure 6. Transcriptional regulation of genes within sugar pathways and those associated with hydrogenosome function for co-culture versus monocultures of *C. churrovis* and *C. churrovis* paired with *M. bryantii*.** The number of genes upregulated (+) or downregulated (-) that are annotated as each enzyme are highlighted in yellow. Enzymes were either identified by Enzyme Commission (EC) number (highlighted in blue) or homology (highlighted in green). Cultures grown on all substrates with the exception of Reed Canary Grass had at least one gene annotated as a pyruvate formate lyase (PFL) upregulated, indicating that coculture with a methanogen may enhance PFL function on certain substrates. PFLs were identified through homology to PFLs identified as hydrogenosome components in the *N. lanati* genome. Select enzymes in a variety of sugar pathways were upregulated in co-culture for some substrates, indicating enhanced production of bottleneck enzymes in sugar pathways. This figure depicts regulation in the cultures grown on Avicel (A) Reed Canary Grass (B), glucose (C), fructose (D), and Xylan (E) substrates.
